# Fear learning sculpts functional brain connectivity at rest beyond the traditional fear network in humans

**DOI:** 10.1101/2020.05.26.115840

**Authors:** Christoph Fraenz, Dorothea Metzen, Christian J. Merz, Helene Selpien, Patrick Friedrich, Sebastian Ocklenburg, Nikolai Axmacher, Erhan Genç

## Abstract

Neuroscientific research has identified specific brain networks involved in the acquisition of fear memories. Using fMRI to assess changes in resting-state functional connectivity (RSFC) induced by fear acquisition, single brain regions from these networks have also been linked to fear memory consolidation. However, previous studies only examined RSFC changes within restricted sets of brain regions or without a proper control group, leaving our knowledge about fear consolidation outside of traditional fear networks incomplete. Here, we tested a group of 84 healthy participants in a differential fear conditioning paradigm and quantified RSFC changes between 358 cortical and 16 subcortical brain areas. Subsequent to fear learning, 21 functional connections exhibited significant RSFC changes. Importantly, these connections were not restricted to the traditional fear networks but also comprised various frontal and visual areas. Our findings indicate that fear memory consolidation is a complex process that integrates relevant information across the entire brain.

## 1. Introduction

The acquisition, maintenance, and extinction of conditioned fear responses are critical functions that help us to evaluate and differentiate potential threats and safety signals in the environment (Craske, Hermans, & Vansteenwegen, 2006; Mineka & Oehlberg, 2008). During the past two decades, the neural circuitry underlying fear learning has been studied extensively in animal models, especially rodents (Milad & Quirk, 2012). Respective efforts were successful in identifying a set of strongly interconnected brain regions involved in the formation of fear memories, namely the hippocampus, specific nuclei of the amygdala, as well as prelimbic and infralimbic cortices (Milad & Quirk, 2012; Orsini & Maren, 2012; Pape & Pare, 2010). Furthermore, they were able to demonstrate that some of these areas exhibit spontaneous reactivations during offline periods subsequent to fear learning, which might indicate their involvement in fear memory consolidation (Pape & Pare, 2010). For example, enhanced hippocampal activity subsequent to fear learning has been observed during sleep (Popa, Duvarci, Popescu, Lena, & Pare, 2010) and awake rest (Carr, Jadhav, & Frank, 2011). In addition, evidence from animal research also suggests that the amygdala and its modulation of synaptic plasticity in other brain regions plays an important role in the consolidation of emotional memories (Roozendaal, McEwen, & Chattarji, 2009). Insights obtained from animal models are commonly used as guiding principles for studies on the acquisition and consolidation of fear memories in the human brain (Milad & Quirk, 2012). However, this approach oftentimes comes with the crucial caveat that large parts of the human brain are not considered for investigation since respective brain areas have not been associated with fear learning in animals before.

In order to study the neural mechanisms underlying the acquisition, maintenance, and extinction of conditioned fear responses in both animals and humans, most studies employ classical Pavlovian fear conditioning paradigms. Usually, these experiments involve the presentation of two neutral stimuli, referred to as conditioned or conditional stimuli. To induce fear learning, one of the conditioned stimuli (CS+) is paired with an aversive stimulus, also called the unconditioned or unconditional stimulus (US), while the other conditioned stimulus (CS−) is never paired with the US. The US is typically administered in the form of electrical stimulation. Subjects learn which of the two conditioned stimuli is followed by the US and show a conditioned fear response (CR). It is common to assess the degree of CR using skin conductance responses (SCRs) or fear ratings (Lonsdorf & Merz, 2017).

The combination of fear acquisition training and neuroimaging methods such as functional magnetic resonance imaging (fMRI) provides an opportunity to reveal specific brain areas involved in fear learning. In humans, respective efforts were able to identify a brain network comprising hippocampus, amygdala, dorsal anterior cingulate cortex (dACC), ventromedial prefrontal cortex (vmPFC), and insula (Milad & Quirk, 2012). To some extent, this so-called “fear network” resembles the set of brain areas that was associated with fear learning in animal studies. However, a large-scale meta-analysis of fMRI studies on fear acquisition in humans identified not one but two complementary networks (Fullana et al., 2016). The first network comprises brain regions that are particularly active during CS+ presentation, which suggests a strong involvement in the processing of potential threat. The respective network strongly resembles what has previously been described as the fear network. In contrast, brain regions from the second network are highly active in response to CS− presentation. Given its vital role in processing non-threatening stimuli, the respective network has been coined the “safety network” (Fullana et al., 2016). It comprises the hippocampus, vmPFC, lateral orbitofrontal cortex, and posterior cingulate cortex (PCC). The authors argued that the processing of the non-threatening CS− is not a passive task but requires emotional and cognitive functions. Hence, they considered the safety network to be as important for the successful evaluation of potentially threatening stimuli as the fear network.

Although fear memories are rapidly acquired (Phelps, 2004), subsequent offline processing, i.e. consolidation of respective memories, is necessary for shaping and maintaining a rich and diverse array of fear related behaviors. One way to examine fear memory consolidation in humans is to assess brain activity while the brain is “at rest”. Again, this can be achieved by neuroimaging methods such as fMRI, which are able to quantify resting-state functional connectivity (RSFC) before and after fear learning. Usually, RSFC is considered to be a valid measure of spontaneous, low frequency fluctuations in brain activity that occur during the absence of sensory input (Fox & Raichle, 2007). However, intrinsic brain activity during periods of rest may not be devoted exclusively to default-mode processing. Alternative perspectives propose that the resting brain actively and selectively processes previous experiences (Miall & Robertson, 2006). Measures of RSFC are relatively stable across time (Birn et al., 2013). However, changes in functional connectivity following behavioral tasks have been observed, especially during immediate post-encoding time periods in which initial stages of memory consolidation are likely to unfold (Hermans et al., 2017). In addition, RSFC patterns observed shortly after a specific task resemble the expression of functional connectivity during that particular task. Such effects have been reported for motor learning (Albert, Robertson, & Miall, 2009; Tung et al., 2013), visual perceptual learning (Guidotti, Del Gratta, Baldassarre, Romani, & Corbetta, 2015; Lewis, Baldassarre, Committeri, Romani, & Corbetta, 2009), attention (Breckel et al., 2013), as well as working memory (Gordon, Breeden, Bean, & Vaidya, 2014) and lexico-semantic cognition (Schlaffke et al., 2017).

Importantly, it has been demonstrated that fear learning processes can alter the RSFC between brain regions from the fear and safety networks. This includes the amygdala, dACC, vmPFC, insula, and hippocampus (P. Feng, Feng, Chen, & Lei, 2014; T. Y. Feng, Feng, & Chen, 2013; Hermans et al., 2017; Schultz, Balderston, & Helmstetter, 2012). However, respective studies leave room for further investigation. First, some studies did not assess RSFC changes in a control group, restricting their analyses to a single within-group comparison (Hermans et al., 2017; Schultz et al., 2012). Thus, the possibility that reported RSFC changes were caused by factors other than fear learning cannot be ruled out. Second, most studies were inspired by animal research and focused their investigations on those brain areas already known to be involved in fear conditioning, leaving thousands of potentially relevant connections untouched. Therefore, it is currently unknown whether fear conditioning tasks induce RSFC changes outside of the fear network. This is particularly problematic as the fear network is not the only neural system associated with the process of differentiating threat and non-threat signals. In addition to the fear and the safety networks, it is possible that further brain areas are involved in fear learning, at least in humans. This is because humans often employ more complex cognitive strategies such as reasoning, decision making, and meta-cognitive judgments on the (assumed) task design in order to resolve even apparently simple paradigms. Reversely, since fear conditioning affects numerous cognitive, emotional and social processes, the fear and safety networks need to interact closely with the brain networks supporting these functions.

In summary, the differentiation between threatening and non-threatening stimuli is likely to rely on at least two distinct neural networks, the fear network and the safety network. As mentioned above, RSFC can be used as a measure to study consolidation processes after fear acquisition. Until now, fMRI research only investigated RSFC changes within the fear network, while largely neglecting the safety network or functional connections outside of the two networks. The current study aimed to examine RSFC changes within the whole brain and tested the hypothesis that fear learning affects RSFC in functional connections outside of the classical fear network, and possibly even beyond the safety network. We investigated RSFC changes between 358 cortical and 16 subcortical brain areas occurring after fear acquisition training. Our main analysis was conducted in three steps and involved a within-group comparison, a between-group comparison, and a correlation analysis that aimed to associate RSFC changes with the degree of CR as quantified by SCRs. We were able to identify RSFC changes in both the fear network and the safety network. Interestingly, we also observed RSFC changes in functional connections that link the fear network to frontal and the safety network to visual areas, suggesting an enhanced exchange of information between these structures during the consolidation of fear memories.

## 2. Material and Methods

### 2.1. Participants

Ninety-five young and healthy participants took part in the current study. They were randomly assigned to an experimental group (*N* = 71), which underwent fear acquisition training with electrical stimulation (see 2.2. Fear Acquisition Paradigm), or a control group (*N* = 24), to which the same stimuli were presented but without electrical stimulation. This was done to examine potential effects of the fear acquisition training by means of between-group analyses (see 2.7. Statistical Analysis). In total, eleven participants from the experimental group had to be excluded due to various reasons. Two participants were removed because of technical difficulties with the electrical stimulation. Three participants were classified as “non-responders” since they did not show valid SCRs in reaction to any of the US presentations (at least 0.05 µS) (Lonsdorf et al., 2019). Six participants reported a CS−/US contingency that was equal to or larger than the reported CS+/US contingency, indicating absent contingency awareness (Tabbert et al., 2011). Hence, all imaging analyses were carried out with data from the remaining 84 participants (54 women).

The age range was 18 to 25 years (*M* = 21.60, *SD* = 1.98). We did not observe significant age differences between male and female participants (*t*(82) = 0.016, *p* = .987). The experimental group included 60 participants (37 women) with a mean age of 21.70 years (*SD* = 1.81) and the control group included 24 participants (17 women) with a mean age of 21.33 years (*SD* = 2.37). Both groups did not differ significantly with regard to age (*t*(82) = 0.766, *p* = .446) or sex distribution (*X*^*2*^(1, *N* = 84) = −0.627, *p* = .428). We also did not observe statistically significant age differences between male and female participants within the experimental (*t*(58) = −0.453, *p* = .653) or control groups (*t*(22) = 0.497, *p* = .624).

In order to control for any potential effects caused by handedness, only right-handed individuals were recruited, as measured by the Edinburgh Handedness Inventory (Oldfield, 1971). All participants had normal or corrected-to-normal vision and were able to understand the instructions given to them orally and in writing. They were either paid for their participation or received course credit. All participants were naive to the purpose of the study and had no former experience with the aversive learning paradigm used for the experiment. Participants reported no history of psychiatric or neurological disorders and matched the standard inclusion criteria for fMRI examinations. The study was approved by the local ethics committee of the Faculty of Psychology at Ruhr University Bochum. All participants provided written informed consent prior to participation and were treated in accordance with the Declaration of Helsinki.

### 2.2. Fear Acquisition Paradigm

While in the fMRI scanner, all participants completed differential fear acquisition training followed by fear extinction training. Importantly, only participants from the experimental group received electrical stimulation during fear acquisition training. In contrast, participants from the control group received no electrical stimulation. Beyond that, participants from both groups underwent the same experimental procedures. Resting-state measurements of around 8 minutes were conducted prior to fear acquisition training, in between both phases, and after fear extinction training. As this study is only concerned with fear acquisition, all procedures related to fear extinction will be reported elsewhere. The stimuli and procedure used for fear acquisition training were modified from (Milad et al., 2007).

After arrival, participants gave written informed consent and filled out a questionnaire on demographic variables. Prior to scanning, all participants were instructed to close their eyes during resting-state scans and to pay close attention to the images being presented during fear acquisition training. They were also told that electrical stimulation may or may not be presented during the experiment. The experimental group and the control group received identical instructions. Electrical stimulation (1 ms pulses with 50 Hz for a duration of 100 ms) was applied as the US using a constant voltage stimulator (STM2000 BIOPAC systems, CA, USA) along with two electrodes attached to the fingertips of the first and second fingers of the right hand. The intensity of electrical stimulation was adjusted for each participant individually prior to the first resting-state scan. For this purpose, electrical stimulation was administered at 30 V and raised in increments of 5 V until participants rated the sensation as very unpleasant but not painful.

During fear acquisition training, a picture of an office room with a switched off desk lamp was used as the context image (Figure 1, top). In each trial, the desk lamp would light up either in blue (CS+) or in yellow (CS−), which served as the two CS. The images were presented using the Presentation software package (Neurobehavioral Systems, Albany, CA) and MR suitable LCD-goggles (Visuastim Digital, Resonance Technology Inc, Northridge, CA). At the beginning of each trial, a white fixation cross was presented on a black background for 6.8 - 9.5 seconds (randomly jittered). Next, the context image was presented for 1 second, which was followed by the presentation of the CS+ or the CS− for another 6 seconds. In the experimental group, the CS+ was paired with electrical stimulation in 62.5% of trials. The stimulation was administered 5.9 seconds after CS+ onset and co-terminated with CS+ offset. Across all 32 trials, CS+ and CS− were presented 16 times each in pseudo-randomized order. The first two trials always consisted of one CS+ and one CS− presentation and so did the last two trials. In the experimental group, the first and last CS+ presentations were always paired with electrical stimulation. There were no trials that presented the same type of CS more than twice in consecutive order. CS+ and CS− presentations were distributed equally across both halves of fear acquisition training.

**Figure 1.**
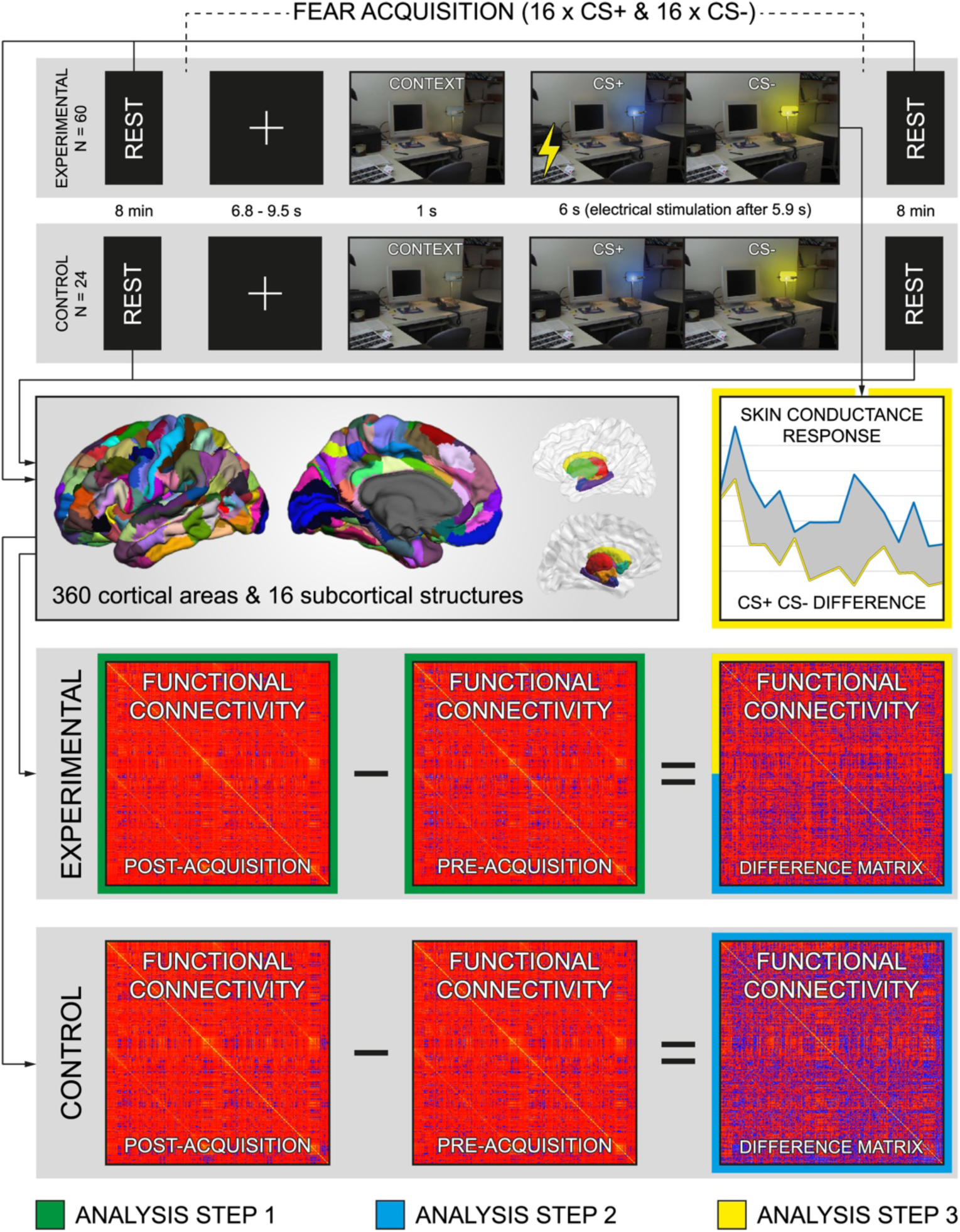
Data acquisition and analysis. The overall sample was split into an experimental (*N* = 60) and a control group (*N* = 24). While in the scanner, both groups participated in fear acquisition training that was preceded and followed by 8-minute-long fMRI resting-state scans. At the beginning of each trial, a white fixation cross was presented on a black background for 6.8 - 9.5 seconds. Next, the context image was presented for 1 second, which was followed by the presentation of the CS+ (blue lamplight) or the CS− (yellow lamplight) for another 6 seconds. In the experimental group, the CS+ was paired with electrical stimulation as unconditioned stimulus (US, indicated by a yellow bolt) in 62.5% of trials administered 5.9 seconds after CS+ onset. The CS+ and CS− were presented 16 times each in pseudo-randomized order. In both groups, brain images obtained from fMRI resting-state scans were parcellated into 360 cortical areas and 16 subcortical structures. The resulting ROIs were subjected to a functional connectivity analysis using BOLD signal correlations. For both groups separately, functional connectivity values from the pre-acquisition matrix were subtracted from the post-acquisition matrix in order to obtain a difference matrix. In the experimental group, fear learning was quantified by subtracting average skin conductance responses to the CS− from those to the CS+. For the first step of the main analysis, indicated by green frames, the pre-acquisition and post-acquisition matrices of the experimental group were compared. Functional connections exhibiting statistically significant differences were subjected to the second step of the main analysis, indicated by light blue frames. Here, the difference matrix of the experimental group was compared to that of the control group. Functional connections demonstrating significant results in both of the aforementioned comparisons were subjected to the third and last step of the main analysis, indicated by yellow frames. Here, conditioned fear response measures quantified via skin conductance responses were regressed on functional connectivity values from the difference matrix of the experimental group.

After fear acquisition training, participants had to rate the contingencies between CS+, CS−, and US. First, they were asked to report the number of electrical stimulations they had received during the experiment. In addition, participants from the experimental group were asked to rate the unpleasantness of the last electrical stimulation on a 9-point Likert scale (1 - “not unpleasant”, 9 - “very unpleasant”) and to report at what rate the blue lamplight and the yellow lamplight had been followed by an electrical stimulation. In contrast, participants from the control group were asked if they had noticed any differences between the two images and how often the blue and the yellow lamplights were presented.

### 2.3. Acquisition of Imaging Data

All imaging data were acquired at the Bergmannsheil hospital in Bochum, Germany, using a 3T Philips Achieva scanner with a 32-channel head coil. Scanning included anatomical imaging, task-based imaging, and resting-state imaging.

#### 2.3.1. Anatomical Imaging

For the purpose of coregistration and brain parcellation, both necessary steps in the connectivity analyses performed in this study, T1-weighted high-resolution anatomical images were acquired (MP-RAGE, TR = 8.2 ms, TE = 3.7 ms, flip angle = 8°, 220 slices, matrix size = 240 mm × 240 mm, resolution = 1 mm × 1 mm × 1 mm). Scanning time was around 6 minutes.

#### 2.3.2. Task-based Imaging

In order to identify specific brain regions showing a significantly pronounced or diminished BOLD response during fear acquisition, we employed echo planar imaging. We obtained a time series of fMRI volumes for each participant from the experimental group while completing fear acquisition training (TR = 2500 ms, TE = 35 ms, flip angle = 90°, 40 slices, matrix size = 112 mm × 112 mm, resolution = 2 mm × 2 mm × 3 mm). The same scanning protocol was used in the control group but obtained imaging data were not used for further analysis. In both cases, scanning time was around 8 minutes.

#### 2.3.3. Resting-state Imaging

For the analysis of functional brain connectivity, fMRI resting-state images were acquired before and after fear acquisition training using echo planar imaging (TR = 2500 ms, TE = 30 ms, flip angle = 90°, 40 slices, matrix size = 112 mm × 112 mm, resolution = 2 mm × 2 mm × 3 mm). Scanning time of each resting-state scan was around 8 minutes.

### 2.4. Acquisition of Skin Conductance Responses

Skin conductance responses were assessed using two Ag/AgCl electrodes filled with isotonic (0.05 NaCl) electrolyte medium placed on the hypothenar eminence right below the fifth finger of the left hand. Data were recorded using Brain Vision Recorder software (Brain Products GmbH, Munich, Germany).

### 2.5. Analysis of Imaging Data

#### 2.5.1. Analysis of Anatomical Data

For the purpose of reconstructing the cortical surfaces of T1-weighted images, we used published surface-based methods in FreeSurfer (http://surfer.nmr.mgh.harvard.edu, version 6.0.0) along with the CBRAIN platform (Sherif et al., 2014). The details of this procedure have been described elsewhere (Dale, Fischl, & Sereno, 1999; Fischl, Sereno, & Dale, 1999). The automatic reconstruction steps included skull stripping, gray and white matter segmentation as well as reconstruction and inflation of the cortical surface. These pre-processing steps were performed for each participant individually. Subsequently, each individual segmentation was quality-controlled slice by slice and inaccuracies of the automatic steps were corrected by manual editing if necessary.

Functional connectivity changes in resting-state data were analyzed between areas defined by the Human Connectome Project’s multi-modal parcellation (HCPMMP) (Glasser et al., 2016) as well as FreeSurfer’s automatic subcortical segmentation (Figure 1, middle). The HCPMMP delineates 180 cortical brain regions per hemisphere and is based on the cortical architecture, function, connectivity and topography from 210 healthy individuals. FreeSurfer’s automatic subcortical segmentation delineates about 40 brain regions, including an array of subcortical structures (thalamus, caudate nucleus, putamen, pallidum, hippocampus, amygdala, accumbens area) and ventricles (lateral ventricle, inferior lateral ventricle, third ventricle, fourth ventricle) as well as the whole cerebral white matter compartment (Fischl et al., 2002). In total, 360 cortical masks, 16 subcortical masks, 8 ventricle masks, and one white matter mask were linearly transformed into the native spaces of the resting-state images that were obtained before and after the acquisition phase. The respective masks served as regions of interest (ROIs) that were used for the functional connectivity analyses.

#### 2.5.2. Analysis of Task-based Data

Task-based data were analyzed by means of FEAT, which is part of the FSL toolbox (http://www.fmrib.ox.ac.uk/fsl, version 6.0.1). Pre-processing of respective images involved motion- and slice-timing correction, spatial smoothing with a 6 mm FWHM Gaussian kernel, high-pass filtering with the cutoff set to 50 seconds, linear registration to the individual’s high-resolution T1-weighted anatomical image, and non-linear registration to the standard stereotaxic space template of the Montreal Neurological Institute (MNI). First-level analyses employed a general linear model (GLM) in order to generate two statistical maps of functional activation (CS+ > CS− and CS− > CS+) including data from all 16 CS+ and 16 CS− presentations. All regressors (context alone, CS+, CS−, US, US omission after CS+ presentation, non-US after CS− presentation) were modeled based on a stick function that was convolved with the canonical hemodynamic response function, without specifically modeling the durations of the different events (i.e. event-related design). Second-level analyses employed random-effects estimation by means of FLAME (FMRIB’s Local Analysis of Mixed Effects). The primary effects of interest included both of the aforementioned contrasts (CS+ > CS− and CS− > CS+) for the whole brain volume. For these contrasts we utilized an FWE-corrected cluster thresholding option, *p*-values < .05, and *Z*-values > 3.1.

#### 2.5.3. Analysis of Resting-state Data

Resting-state data were pre-processed using MELODIC, which is also part of the FSL toolbox. Images were pre-processed in a number of steps: Discarding the first two EPI volumes from each resting-state scan to allow for signal equilibration, motion and slice-timing correction, high-pass temporal frequency filtering (0.005 Hz). Spatial smoothing was not applied in order to avoid the introduction of spurious correlations in neighboring voxels. For each region of interest, we calculated a mean resting-state time course by averaging the pre-processed time courses of corresponding voxels. Two ROIs from the PCC, labeled “L_23c” and “R_23c” by the HCPMMP, had to be excluded from our analyses since their average time courses turned out to be 0 in every participant.

We computed partial correlations between the average time courses of the remaining 358 cortical and 16 subcortical regions, while controlling for several nuisance variables. We regressed out the trajectories of 6 head motion parameters as well as the mean time courses extracted from the white matter and ventricle masks (Genc, Schoelvinck, Bergmann, Singer, & Kohler, 2016). The resulting correlation coefficients were subjected to a Fisher *z*-transformation (Fisher, 1921) in order to receive normally distributed data suitable for further testing. By following this approach, we obtained two symmetrical 374-by-374 matrices with data assessed before (Supplementary Figure 1 and Supplementary Figure 2) and after fear acquisition training (Supplementary Figure 3 and Supplementary Figure 4), respectively (Figure 1, bottom). Each cell of the two matrices contained a Fisher transformed correlation coefficient as a measure of functional connectivity between a specific pair of ROIs. When accounting for the symmetry of both matrices and neglecting the self-correlations on their diagonals, each matrix comprised 69751 individual connections. For each participant we calculated the difference between both matrices by subtracting the pre-acquisition matrix from the post-acquisition matrix (Supplementary Figure 5 and Supplementary Figure 6).

### 2.6. Analysis of Skin Conductance Responses

Raw SCR data from the experimental group were pre-processed with Brain Vision Analyzer software (Brain Products GmbH, Munich, Germany). All further analyses were conducted semi-automatically using MATLAB, version 9.6.0.1114505 (R2019a, The MathWorks Inc., Natrick, MA). SCR to CS presentation was defined as the maximum amplitude recorded within the time window starting 1 s after CS onset and ending 6.5 s after CS onset. The CR was quantified as the difference between the average SCR across CS+ trials and the average SCR across CS− trials (Figure 1, middle). SCR to US presentation was defined as the maximum amplitude recorded within the time window starting 6.5 s after CS onset and ending 12 s after CS onset. The unconditioned response (UR) was quantified as the difference between the average SCR across CS+ trials with electrical stimulation and the average SCR across CS− trials. In three cases, we were unable to compute measures of CR since data on SCRs to either CS+ or CS− were missing. Therefore, the final step our analysis, which involved regressing SCR measures of CR on RSFC changes, had to be carried out on data from the remaining 57 participants in the experimental group (Figure 2).

**Figure 2.**
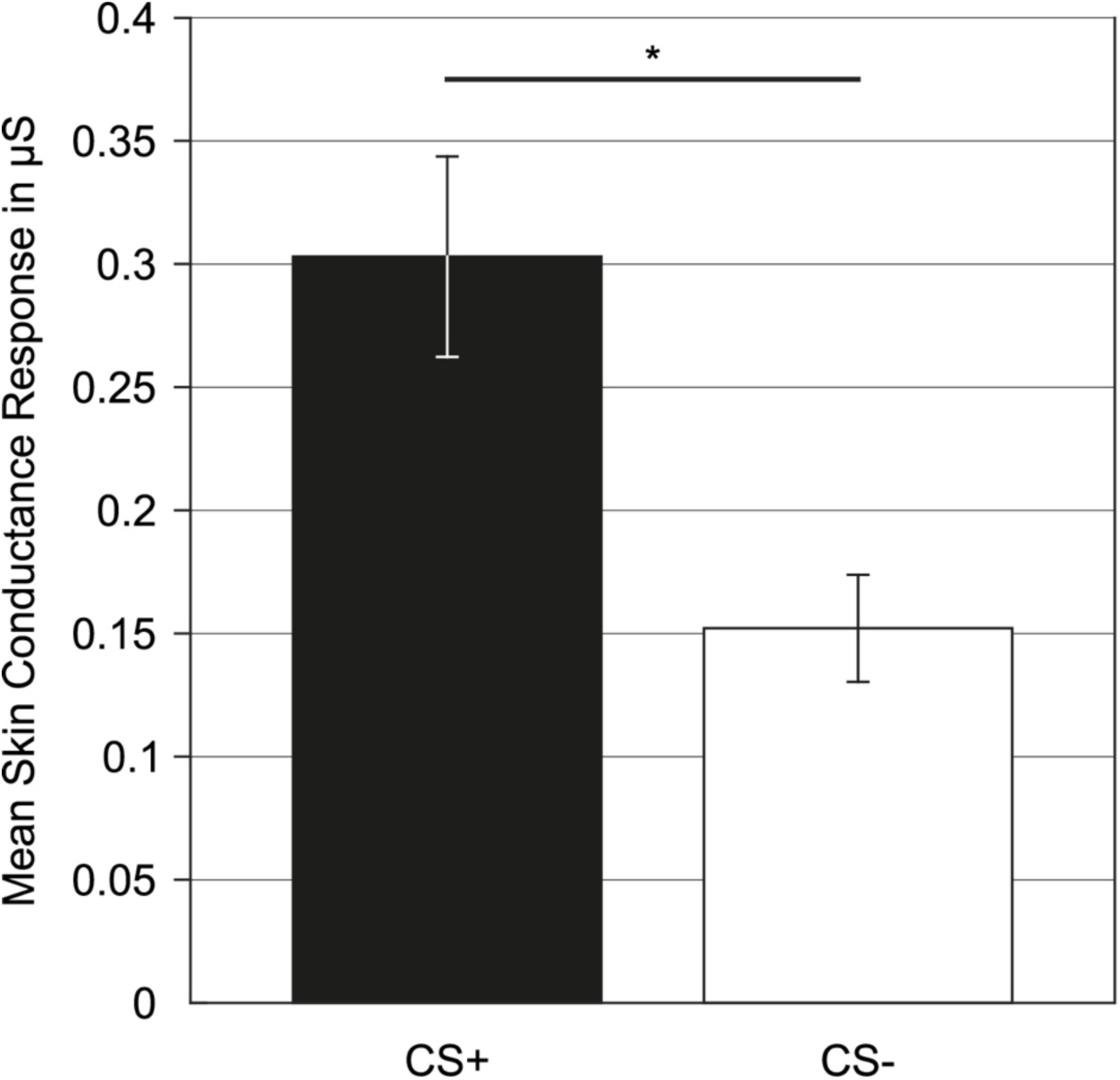
Mean skin conductance responses. Bar plots showing the mean skin conductance responses in reaction to CS+ (black bar) and CS− (white bar) presentations in microSiemens (µS). Data were averaged across all 16 CS+ and 16 CS− trials, respectively. Error bars represent standard errors (* *p* < .001).

### 2.7. Statistical Analysis

All statistical analyses were carried out using MATLAB, version 9.3.0.713579 (R2017b, The MathWorks Inc., Natick, MA). For all analyses, we employed linear parametric methods. Testing was two-tailed with an α-level of .05, which was corrected for multiple comparisons using the Benjamini-Hochberg method (Benjamini & Hochberg, 1995).

For the purpose of investigating fear memory consolidation, we examined learning-induced changes in RSFC. The respective analysis was carried out sequentially and comprised three major steps. First, we performed paired-sample *t*-tests between functional connectivity values from the pre-acquisition and the post-acquisition matrices. Here, our goal was to assess which of the 69751 functional connections showed significant increases or decreases in their Fisher transformed BOLD signal correlations after fear acquisition training. Hence, respective tests were carried out for the experimental group exclusively and corrected α-levels were in the range between .05 / 69751 = .000007 and .05 as defined by the Benjamini-Hochberg method. Second, all functional connections reaching statistical significance in the first step were subjected to a comparison between the experimental and control group. For this purpose, the difference matrices (see 2.5.3. Analysis of Resting-state Data) from the experimental group were compared to those from the control group using two-sample *t*-tests. Again, we accounted for multiple comparisons using the Benjamini-Hochberg method. This time, the α-levels were corrected according to the number of functional connections that exhibited statistical significance after the first step of our analysis.

Third, all remaining functional connections, showing both significant changes between the pre-acquisition and post-acquisition matrices as well as between the experimental and control groups, were considered for the final step of our analysis. Here we computed a hierarchical linear regression analysis with RSFC changes serving as independent variables and CR as quantified by SCRs serving as the dependent variable (see 2.6. Analysis of Skin Conductance Responses). We employed a backward elimination approach with a threshold of *p* = .05 to identify significant predictors of CR. Initially, this method places all potential predictors in a regression model and computes their individual contributions towards the dependent variable. With every step of the analysis, the predictor with the least contribution in terms of the *p*-value is excluded from the model. The individual contributions of the remaining predictors are then re-estimated in an iterative manner. Elimination stops once the *p*-values of all remaining predictors are below the desired threshold, in this case *p* = .05..

## 3. Results

### 3.1. Fear Memory Acquisition

Participants showed significantly greater SCRs in reaction to CS+ presentations relative to CS− presentations (*t*(56) = 5.465, *p* < .001) (Figure 2), indicating successful fear learning. Furthermore, fMRI data obtained during fear acquisition training revealed distinct patterns of functional activation in reaction to CS+ and CS− presentations. Overall, our data were in good accordance with the results of a large-scale meta-analysis on the functional correlates of fear acquisition (Fullana et al., 2016). Voxel clusters exhibiting significant (*p* < .05, *Z* > 3.1, FWE-corrected) BOLD signal contrasts in our data either overlapped with clusters from the fear and safety networks or were directly adjacent to them (Supplementary Figure 7 and Supplementary Figure 8). With regard to the fear network, we were able to replicate relevant structures like the dACC, thalamus, and secondary somatosensory cortex in the contrast CS+ minus CS−. Likewise, relevant structures from the safety network, namely the angular gyrus, primary somatosensory cortex, parahippocampal gyrus, PCC, vmPFC, and lateral orbitofrontal cortex, could also be identified in our data in the contrast CS− minus CS+.

### 3.2. Fear Memory Consolidation

For the first step of our analysis, we employed paired-sample *t*-tests to compare the experimental group’s pre-acquisition matrices (Supplementary Figure 1) to its post-acquisition matrices (Supplementary Figure 3). In total, we observed 2026 functional connections that exhibited significant changes after correction for multiple comparisons (see 2.7. Statistical Analysis). It is noteworthy that 2009 out of these 2026 significant connections (99.16%) showed increases in their Fisher transformed BOLD signal correlations, meaning that positive coefficients became larger, negative coefficients became smaller, or negative coefficients became positive. This pronounced shift towards more positive coefficients was not only present among the 2026 significant connections but could also be observed for the overall matrix of functional connections. Here, 53974 out of 69751 connections (77.38%) showed increases, while only 15777 (22.62%) showed decreases. We observed a similar, albeit significantly less pronounced (*X*^*2*^(1, *N* = 139502) = 3348.12, *p* < .001), pattern in the control group with 44099 connections (63.22%) showing increases and 25652 connections (36.78%) showing decreases.

For the second step of our analysis, all functional connections that exhibited statistically significant changes between the pre-acquisition and post-acquisition matrices were subjected to a comparison between the experimental and control groups. For this purpose, we compared the difference matrices of the two groups (Supplementary Figure 5 and Supplementary Figure 6) with each other using two-sample *t*-tests. Again, the α-levels were corrected using the Benjamini-Hochberg method and ranged from .05 / 2026 = .00002 to .05 given the number of functional connections (2026) that were considered in this step. As a result, we identified 21 functional connections for which the experimental group demonstrated RSFC changes that were significantly different from those shown by the control group (Supplementary Table 1). Interestingly, all changes within the experimental group were increases, whereas the control group only exhibited decreases. Respective functional connections were almost exclusively constituted by cortical areas. The connection comprising the subcortical ROI “lh_caudate” and the cortical ROI “L_45” (corresponding to the caudate nucleus and Brodmann area 45) was the only exception. There were 17 interhemispheric and only 4 intrahemispheric connections, namely ROI pairs “L_AVI” and “L_a10p” (corresponding to the anterior insular cortex and a subregion of Brodmann area 10), “lh_caudate” and “L_45”, “R_v23ab” and “R_PH” (corresponding to a subregion of Brodmann area 23 and a subregion of the visual cortex), as well as “L_IFSa” and “L_a10p” (corresponding to the anterior extent of the inferior frontal sulcus and a subregion of Brodmann area 10) (Figure 3). Changes in RSFC ranged from 0.0555 to 0.1037 in the experimental group and from −0.0297 to −0.0935 in the control group. In the experimental group, the functional connection with the highest increase (0.1037) was between the ROIs “L_AVI” and “L_a10p”. The same functional connection exhibited a decrease (−0.0546) in the control group (*t*(82) = 3.7150, *p* = .00004).

**Figure 3.**
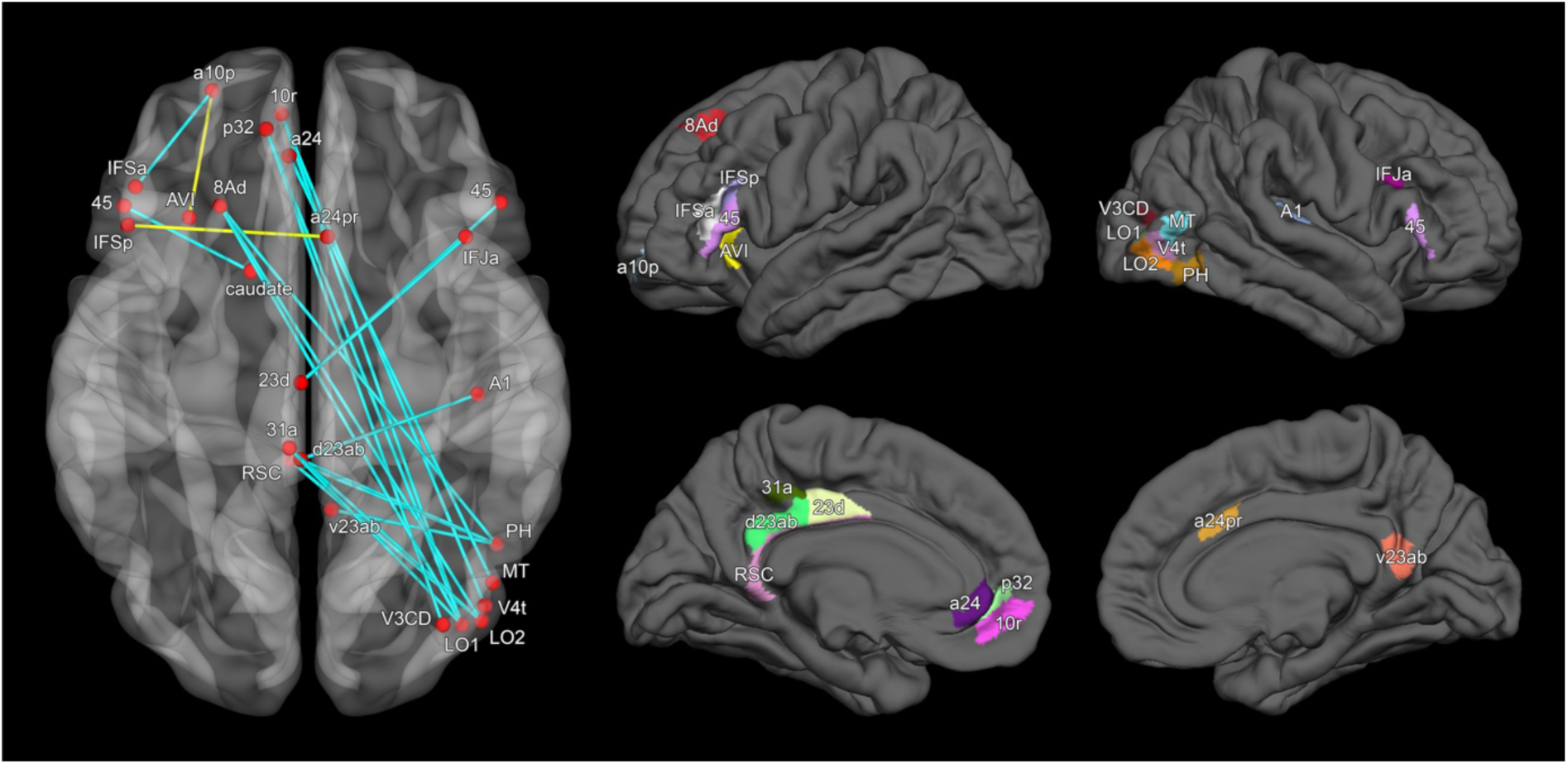
Learning-related changes in functional connectivity. On the left-hand side, functional connections are shown as light blue or yellow lines connecting pairs of brain areas, which are depicted as red nodes on a semi-transparent brain. In the first step of the main analysis, all functional connections showing significant increases or decreases in response to fear acquisition training were identified. In the second step, respective connectivity changes were subjected to a comparison between the experimental and control groups. In the third and final step, a multiple regression analysis with backward elimination was used to identify functional connections in which significant functional connectivity changes were correlated with conditioned fear response measures quantified via skin conductance responses. Functional connections which exhibited statistically significant results in both of the first two steps are depicted in light blue. Functional connections which reached statistical significance in all three steps are depicted in yellow. On the right-hand side, all brain areas constituting statistically significant connections are shown on lateral and medial views of cortical surfaces. Naming of brain areas is based on the Human Connectome Project’s Multimodal Parcellation as well as FreeSurfer’s Aseg Atlas.

For the third step of our analysis, we computed a hierarchical linear regression analysis with a backward elimination approach. The RSFC changes of the remaining 21 connections served as independent variables, while CR quantified via SCR measures obtained during fear acquisition training served as the dependent variable (see 2.6. Analysis of Skin Conductance Responses). With all variables included, the regression model was not significant for predicting CR (*R*^*2*^ = 0.416, *R*^*2*^_Adjusted_ = 0.065, *F*(21, 35) = 1.187, *p* = 0.319). The step-wise elimination of variables yielded a regression model with two functional connections remaining as significant predictors (*R*^*2*^ = 0.157, *R*^*2*^_Adjusted_ = 0.125, *F*(2, 54) = 5.010, *p* = 0.010). Specifically, these were the functional connections between HCPMMP areas “R_a24pr” and “L_IFSp” (corresponding to a subregion of Brodmann area 24 and the posterior extent of the inferior frontal sulcus) (*B* = 0.277, *p* = .038) as well as between “L_AVI” and “L_a10p” (*B* = −0.368, *p* = .006) (Supplementary Figure 9). The results from all three steps of our analysis, along with a visualization of HCPMMP areas, are shown in Figure 3, Supplementary Figure 10, and Supplementary Table 1.

## 4. Discussion

The primary goal of this study was to investigate the association between fear acquisition and subsequent RSFC changes. In particular, we were interested in respective changes exhibited by functional connections outside of brain networks that have previously been linked to fear processing. To this end, we followed a whole-brain approach and examined RSFC changes between 358 cortical and 16 subcortical areas in a sample of 84 healthy participants.

In comparison to previous studies on the association between RSFC changes and fear acquisition (P. Feng et al., 2014; T. Y. Feng et al., 2013; Hermans et al., 2017; Schultz et al., 2012), our approach differs in two crucial aspects. First, we did not restrict our analysis to a preselected set of ROIs from the fear or safety networks but comprehensively investigated RSFC changes across the entire brain, namely in almost 70000 functional connections. Second, in order to ensure the specificity of these effects, we compared the RSFC changes found in the experimental group with those found in a control group. Our results clearly demonstrate that lack of a control group may result in unspecific effects: The analysis of our experimental group yielded significant RSFC changes in over 2000 different functional connections. However, when conducting a comparison between the experimental and control groups, only 21 of these changes in connectivity were specific to the experimental group. In other words, about 99% of RSFC changes observed within the experimental group could not be attributed to fear learning itself. For this reason, well-controlled designs are crucial for understanding the neural mechanisms underlying fear memory consolidation. It has to be noted that some of the previously conducted studies have used such between-group designs (P. Feng et al., 2014; T. Y. Feng et al., 2013). However, none of them investigated RSFC changes outside of the fear or safety networks. This is an important aspect as indicated by the three types of functional connections for which we were able to observe significant RSFC changes. First, there were connections constituted by brain areas from the fear and safety networks. Second, we also found functional connections linking the safety network to several right-hemispheric visual areas. Third, we could identify connections running between the fear network and frontal areas, predominantly in the left hemisphere. All of the above will be discussed in the following. In addition to the aforementioned overlap in task-related data (see 3.1. Fear Memory Acquisition), functional connections exhibiting significant RSFC changes after fear acquisition training (Figure 3) also matched the structure of the fear and safety networks. In more detail, we identified five functional connections that were, at least in part, constituted by brain areas from the fear network. Among these were the left caudate nucleus as well as HCPMMP areas “L_AVI”, “L_45”, “L_23d”, and “R_a24pr”, which correspond to the anterior insular cortex (AIC), Brodmann area 45, and dorsal compartments of the cingulate cortex. The involvement of AIC and dACC is especially interesting since joint activation of these “cingulofrontal cortex” structures is a common observation made in many human fMRI studies (Mechias, Etkin, & Kalisch, 2010; Sehlmeyer et al., 2009). It has been proposed that both structures serve as key components of an autonomic-interoceptive network. Within this framework, the AIC is conceptualized to generate an integrated representation of the body’s cognitive, affective, and physical state, while the dACC is thought to be an output region that initiates homeostatic autonomic and behavioral responses (Medford & Critchley, 2010). As shown by (Fullana et al., 2016), the ventral AIC and rostral dACC, among other regions, exhibit fear-conditioned brain activation even when confounding effects caused by the US are controlled. From this, it can be concluded that functional activation of these brain regions in response to CS+ presentation is likely to be “purely anticipatory in nature and unrelated to the generation of defensive autonomic responses to the US” (Fullana et al., 2016). It is conceivable that both regions are involved in generating mental representations of the body’s inner state after experiencing the US. Moreover, both regions might maintain such representations over the course of the complete fear acquisition training. With each trial, additional visual information about the co-occurrence of US and CS might be integrated with the initially generated body representation to form a stable trace in fear memory. Importantly, the rostral dACC and ventral AIC closely resemble the HCPMMP areas “R_a24pr” and “L_AVI” that we could identify in our analysis. Hence, RSFC changes between these regions may represent a form of consolidation initiated by the mechanisms of fear learning described above.

More than half of the functional connections exhibiting significant RSFC changes were constituted by brain areas from the safety network. Except for HCPMMP area “L_8Ad”, located along the superior frontal gyrus, all of these regions, namely “L_d23ab, ‘‘L_10r”, “L_a24”, “L_p32”, “L_31a”, and “L_RSC”, fall onto the brain’s mid-sagittal plane. To be exact, HCPMMP areas “L_a24” and “L_p32” do not directly overlap with parts of the safety network but together with area “L_10r” they form a cluster in the vmPFC closely resembling one of the safety network’s main components. Moreover, a large portion of area “L_RSC” does overlap with a cluster of the safety network in the retrosplenial cortex but due to “L_RSC” extending across the whole posterior half of the cingulum, it also overlaps with a cluster from the fear network.

In general, the structure of the safety network shows some overlap with the default-mode network, which is active during phases requiring little or no cognitive effort (Raichle, 2015). Despite the close resemblance between the safety and default-mode networks, the former is likely to reflect more than task inactivity given that CS− processing demands attention and evaluative processes (Harrison et al., 2017). For example, several safety network components like the vmPFC, PCC, and hippocampus have been reported to show functional activation while participants imagined pleasant scenarios (Leknes, Lee, Berna, Andersson, & Tracey, 2011).

Moreover, the vmPFC has been hypothesized to serve an important role in differentiating between CS+ and CS− by exerting fear response inhibition. It is thought to encode the intrinsic reward value of threatening and non-threatening stimuli. Therefore, functional activation in response to CS− presentations might be the result of vmPFC processing safety signals with high reward values indicating the omission of subsequent US presentation (Harrison et al., 2017). Brain regions from the PCC and retrosplenial cortex have been proposed to form an extended episodic memory network together with other structures such as the hippocampus. Functional co-activation of these regions in response to the CS− might serve the general purpose of encoding the association between CS and US (Holt, Coombs, Zeidan, Goff, & Milad, 2012). Given these mechanisms exerted by the safety network, it is reasonable to assume that RSFC changes in functional connections emerging from HCPMMP areas in the vmPFC (‘‘L_10r”, “L_a24”, “L_p32”) or the PCC (“L_d23ab, “L_RSC”) represent consolidation processes strengthening the perception of CS− as a safety signal.

In addition to RSFC changes within the fear and safety networks, we also identified RSFC changes in functional connections linking both networks to other areas of the brain. More specifically, we observed enhanced functional connections between components of the safety network and a right-hemispheric cluster of visual brain regions, namely HCPMMP areas “R_LO1”, “R_LO2”, “R_PH”, and “R_V4t”. Respective areas extend from the lateral occipital cortex to the inferior temporal gyrus and are associated with the processing of color and form (Gegenfurtner, 2003; Goddard, 2017; Lafer-Sousa, Conway, & Kanwisher, 2016). To our knowledge, these specific brain regions have not been discussed in previous research on fear acquisition let alone fear consolidation. However, an involvement of primary visual cortex (V1) has been demonstrated in both primates (Li, Yan, Guo, & Li, 2019) and humans (Knight, Smith, Stein, & Helmstetter, 1999; Shalev, Paz, & Avidan, 2018; Stolarova, Keil, & Moratti, 2006). The majority of these studies used visual grating stimuli of different orientation as CS and assessed functional activity in V1 by means of electrophysiology or fMRI. The data revealed distinct patterns of activity in response to CS+ and CS−. Since the orientation of visually presented gratings is known to be processed by V1, it is plausible that V1 is also involved in fear learning paradigms employing such stimuli. Concordantly, a study employing fear acquisition training with face stimuli as CS was able to demonstrate enhanced RSFC in functional connections emerging from the fear network and the fusiform face area (Hermans et al., 2017). Thus, sensory areas dedicated to the processing of a specific CS type appear to play an important role in both the acquisition and consolidation of respective fear memory traces. Our results are in good accordance with these observations. Brain regions from the right visual cortex demonstrated increased functional activation during fear learning (Supplementary Figure 7 and Supplementary Figure 8) and functional connections emerging from said regions showed enhanced RSFC shortly thereafter. Since the color of our CS served as the primary indicator of threat or safety, it is plausible that fear memory consolidation relies on increased communication between safety network components and visual brain areas processing color and form. This insight has not been revealed by previous studies concerned with RSFC changes subsequent to fear acquisition as their selection of ROIs comprised the fear and safety networks but did not include visual areas.

It is particularly striking that the occipito-temporal brain areas formed a cluster only in the right but not in the left hemisphere. This observation is well in line with previous findings on hemispheric asymmetries in fear processing. Evidence from a lesion study in human patients with focal brain damage suggests that visual processing of negative emotions such as fear and sadness is particularly impaired after damage to the right inferior parietal cortex and in the right mesial anterior infracalcarine cortex (Adolphs, Damasio, Tranel, & Damasio, 1996). A similar relationship was not found for the left infracalcerine cortex, suggesting a specific role of the right occipital cortex in visual fear processing.

In view of the fear network, we observed enhanced functional connections to several frontal brain regions, namely HCPMMP areas “L_IFSa”, “L_IFSp”, “R_IFJa”, and “L_a10p”. Here, the connection between “L_IFSp” and “R_a24pr”, corresponding to the inferior frontal sulcus and dACC, as well as the connection between “L_a10p” and “L_AVI”, corresponding to Brodmann area 10 and the AIC, are of particular interest. In both cases, changes in RSFC were associated with the degree of CR as measured by SCRs.

The functional connection between “L_IFSp” and “R_a24pr” exhibited a positive correlation between RSFC changes and CR. This implies that participants with a stronger increase in RSFC demonstrated a more distinct reaction towards the presentation of threatening (CS+) and non-threatening stimuli (CS−). As outlined above, HCPMMP area “R_a24pr”, which corresponds to the rostral segment of the dACC, is a key component of the fear network (Fullana et al., 2016). Furthermore, it has been proposed to form an autonomic-interoceptive network together with other brain regions such as the AIC (Medford & Critchley, 2010). In this regard, “R_a24pr” is considered to initiate homeostatic autonomic and behavioral responses in reaction to mental representations of the body’s inner state provided by the AIC. Interestingly, HCPMMP area “L_IFSp” is located directly adjacent to the inferior frontal gyrus (IFG). The IFG has been associated with the selection of competing mental representations (Thompson-Schill, D’Esposito, Aguirre, & Farah, 1997; Zhang, Feng, Fox, Gao, & Tan, 2004), especially under non-response conditions that do not require the respective selection to be verbalized (Milham et al., 2001). In addition, the IFG was also found to exhibit increased functional activity during tasks that tap into inhibitory processes (D’Esposito, Postle, Jonides, & Smith, 1999). These findings are in good accordance with the results of the current study since both selection and inhibition are vital aspects of fear learning. It is conceivable that the AIC provides HCPMMP area “L_IFSp” with mental representations of the body’s inner state after US presentation or US omission. With each trial of fear acquisition training, area “L_IFSp” might select one of these representations and inhibit the other, depending on the type of CS being presented, with CS+ indicating threat and CS− indicating safety. The selected mental representation might then be processed by the rostral dACC (“R_a24pr”) in order to generate an adequate response. Therefore, increased interaction between these regions is likely to enhance the differentiation of CS+ and CS−. This mechanism, in turn, might underly our observation that increases in RSFC between “L_IFSp” and “R_a24pr” are positively associated with CR as quantified by SCRs.

The functional connection between “L_a10p” and “L_AVI” exhibited a negative correlation between RSFC changes and CR. This implies that participants with a weaker increase in RSFC demonstrated a more distinct reaction towards the presentation of threatening (CS+) and non-threatening stimuli (CS−). There is a large body of research demonstrating the existence of anatomical connections between Brodmann area 10 and the AIC (Peng, Steele, Becerra, & Borsook, 2018; Öngür, Ferry, & Price, 2003). With regard to functional activation, both areas have been associated with processing pain, pain anticipation, and pain memory (Peng et al., 2018). Results on the functional interplay between Brodmann area 10 and the AIC have been reported by a study investigating physical stress responses to trauma-related stimuli in war veterans. Strong reactions were associated with functional deactivation of Brodmann area 10 and functional activation of the insula, while weak reactions were associated with the opposite pattern (King et al., 2009). Based on this observation, the authors proposed that Brodmann area 10 might have an inhibitory effect on fear responses towards threatening stimuli. This hypothesis is in accordance with our data. As demonstrated by the meta-analysis of (Fullana et al., 2016), the AIC is a key component of the fear network, showing functional activation in response to threatening stimuli (CS+). Co-activation of Brodmann area 10, or one of its subregions like HCPMMP area “L_a10p”, might indicate some form of inhibition being exerted on the AIC. This could lead to a weaker SCR towards CS+ presentation and, in turn, a less pronounced differentiation between CS+ and CS−. Therefore, fear response inhibition caused by co-activation of “L_a10p” and “L_AVI” might produce a shift in RSFC towards more positive coefficients. On the contrary, participants exhibiting a shift towards more negative RSFC coefficients as well as a more pronounced CR might do so because of disentangled functional activation of areas “L_a10p” and “L_AVI” during fear acquisition, possibly indicating the absence of fear response inhibition. It is also noteworthy that the modulation of functional connectivity between left frontal areas subsequent to fear conditioning is well in line with an established model of hemispheric asymmetries in emotion processing and emotion regulation. According to the asymmetric inhibition model (Grimshaw & Carmel, 2014), left-lateralized executive control processes inhibit negative emotional distractors. This supports the idea that the pattern observed in the present data reflects a form of emotion regulation. While “L_AVI” is involved in the processing of specific fear-related stimuli presented during fear acquisition training, “L_a10p” is likely to regulate the emotional reaction to these stimuli. In summary, we were able to confirm previous observations suggesting an involvement of fear network components in the consolidation of fear memories. Moreover, we observed the same for brain areas from the safety network, which - until now - has been widely neglected by studies on the association between fear acquisition and RSFC changes. Furthermore, we could show that significant RSFC changes subsequent to fear acquisition also comprise brain areas beyond known fear and safety network components. The safety network exhibited enhanced functional connections with visual areas encoding CS specific information, namely color and form, whereas the fear network demonstrated increased RSFC with several frontal areas. Interestingly, we found RSFC changes of two functional connections from the frontal cortex to be correlated with SCR measures of CR. The functional nature of the brain areas involved in these connections suggests that processes of selection and inhibition play an important role for differentiating between threatening and non-threatening stimuli.

## 5. Conclusions

Our findings demonstrate that fear acquisition and subsequent fear consolidation are accompanied by a wide range of complex functional interactions between several brain regions. Importantly, these regions are not exclusively found in previously proposed brain networks associated with the processing of threatening and non-threatening stimuli, but also include visual and frontal areas. On a methodological level, our results also highlight the importance of proper control group designs as the majority of observed RSFC changes in the experimental group disappeared after comparing them to a respective control group. Moreover, our findings advocate the idea of following a whole-brain approach when investigating the neural basis of fear learning and consolidation.

## 6. Acknowledgements

This work is part of the SFB 1280 projects A02, A03, A09, and F02 and was supported by the Deutsche Forschungsgemeinschaft (DFG) (project number 31680338). The authors thank M. A. Fullana and B. J. Harrison for providing us with the original maps of functional brain activation from their meta-analysis on human fear conditioning. Further, the authors thank PHILIPS Germany (Burkhard Mädler) for scientific support with the MRI measurements as well as Tobias Otto for technical support. Finally, the authors would like to thank all research assistants for their support with data acquisition.

