## Supplementary Figures 1 - 10 for "Fear learning sculpts functional brain connectivity at rest beyond the traditional fear network in humans"

### Supplementary Figure 1

Functional brain connectivity assessed in the experimental group prior to fear acquisition training

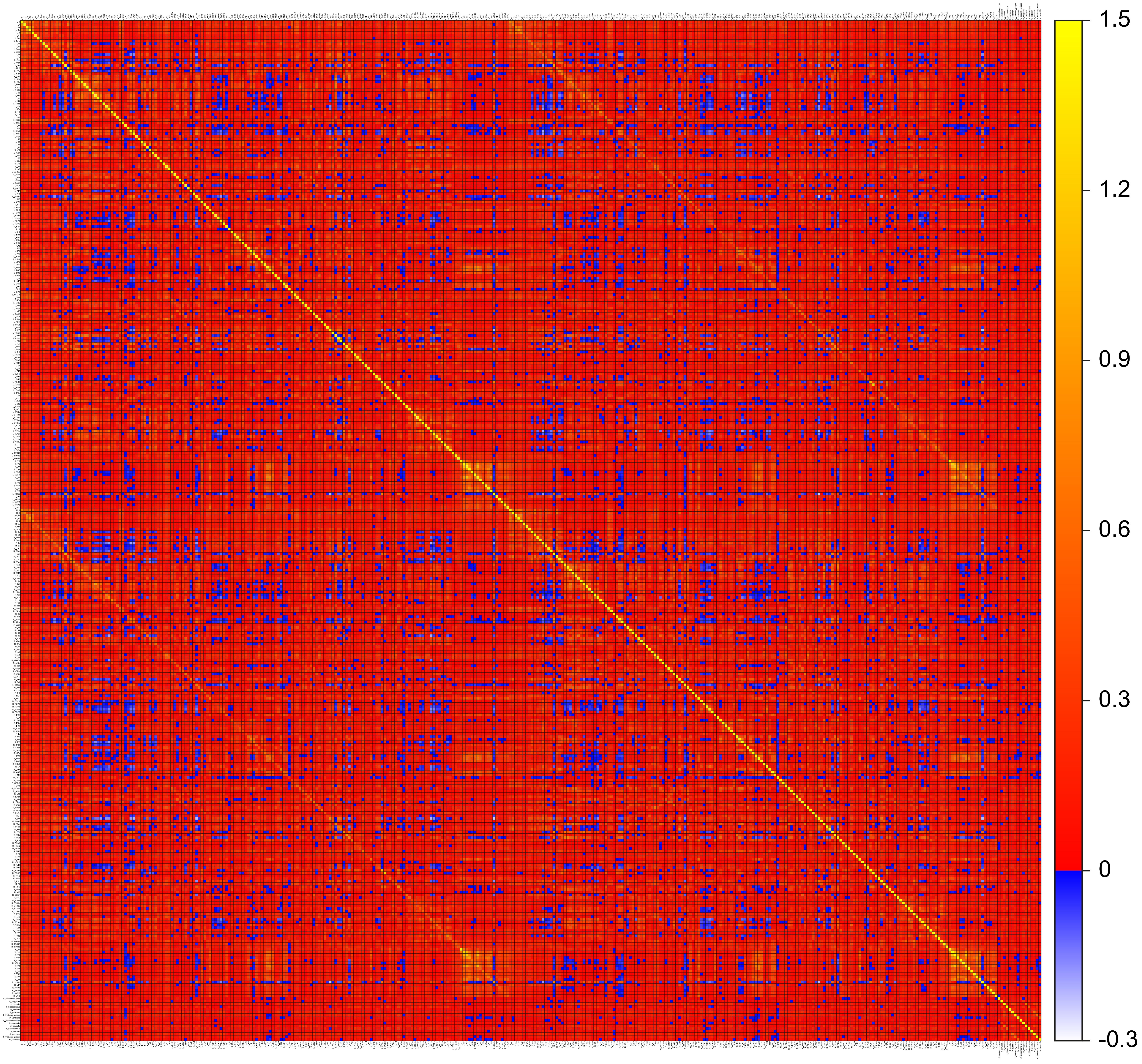

### Supplementary Figure 2

Functional brain connectivity assessed in the control group prior to fear acquisition training

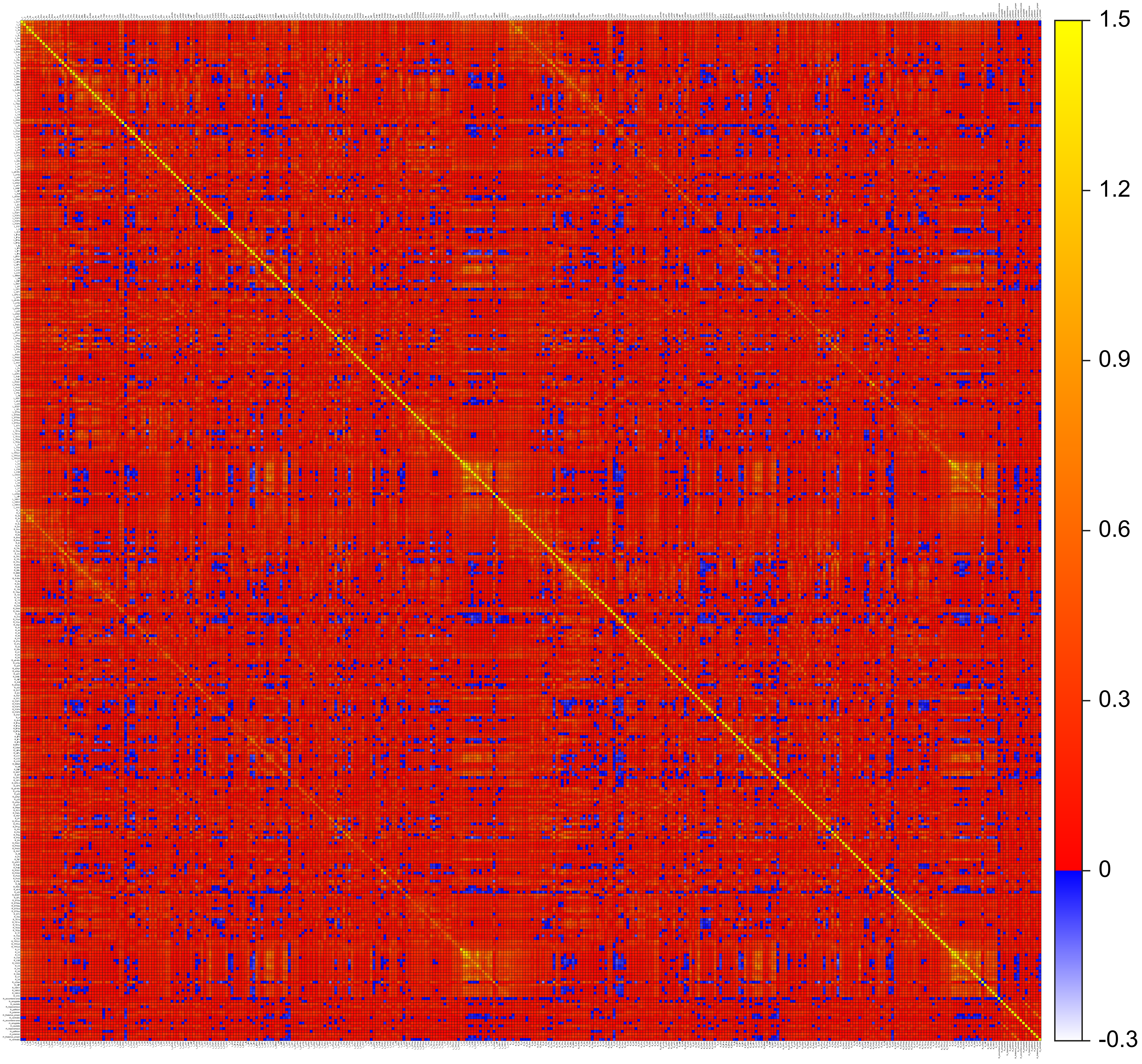

### Supplementary Figure 3

Functional brain connectivity assessed in the experimental group after fear acquisition training

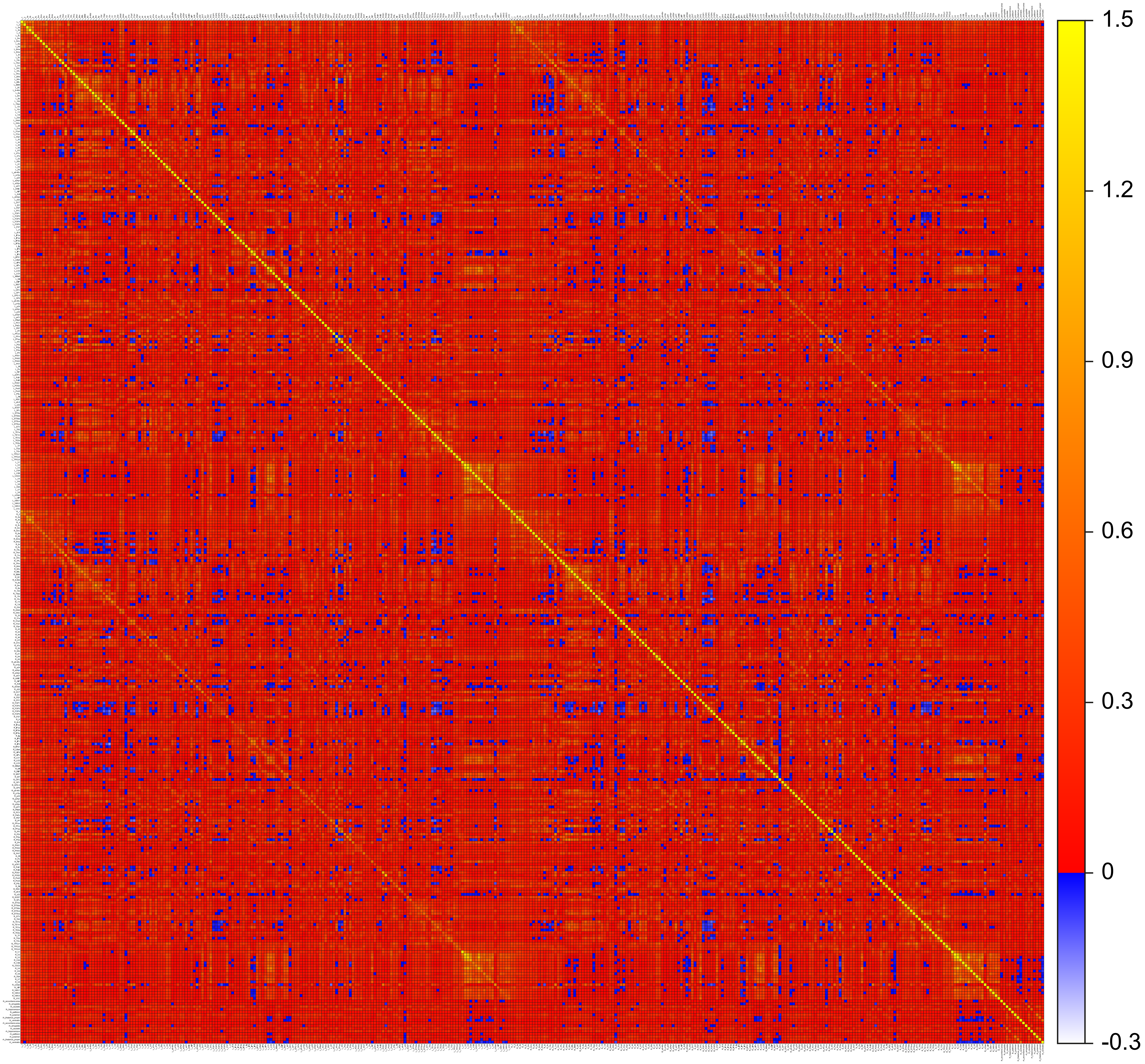

### Supplementary Figure 4

Functional brain connectivity assessed in the control group after fear acquisition training

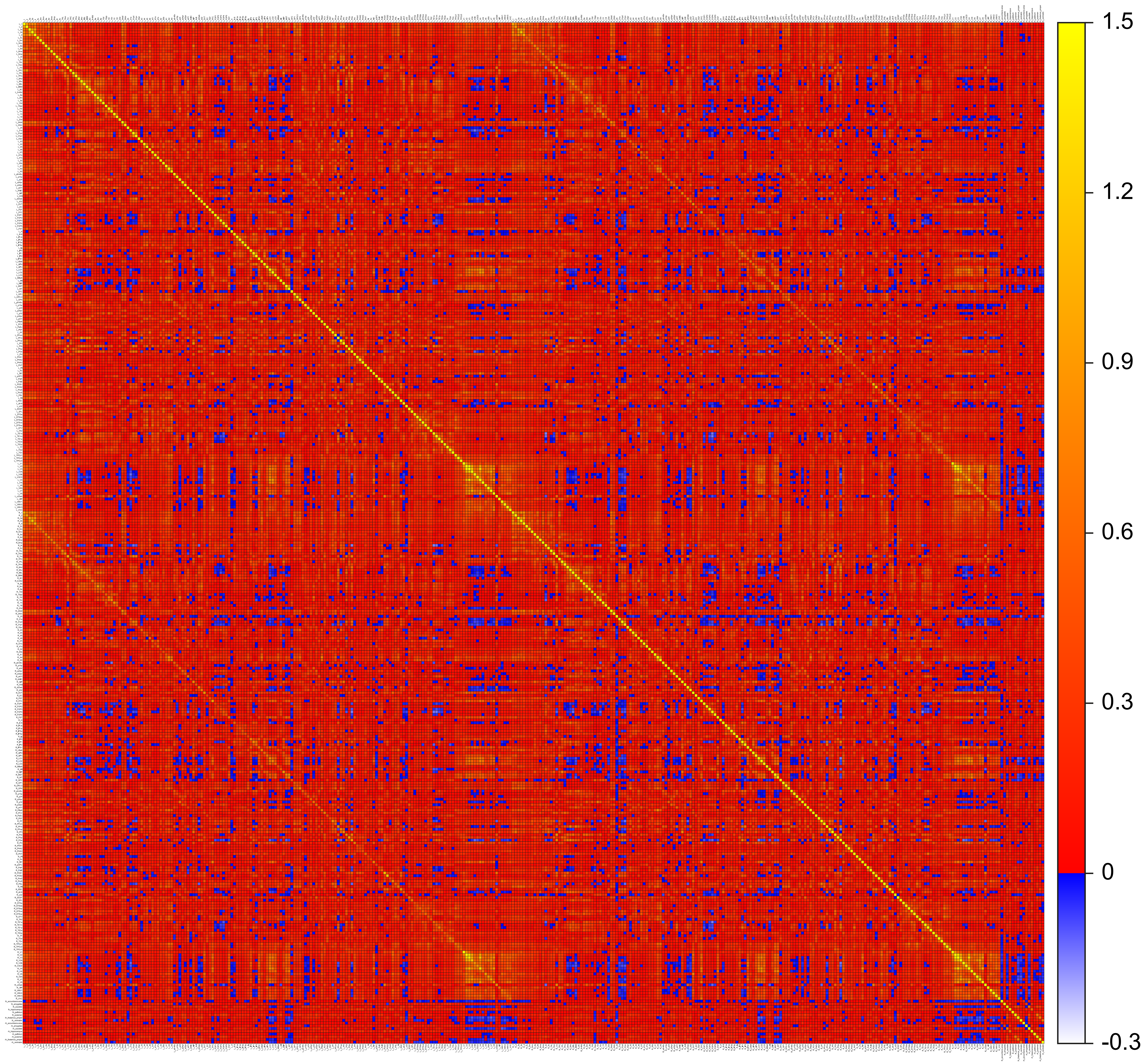

### Supplementary Figure 5

Changes in functional brain connectivity observed in the experimental group after fear acquisition training

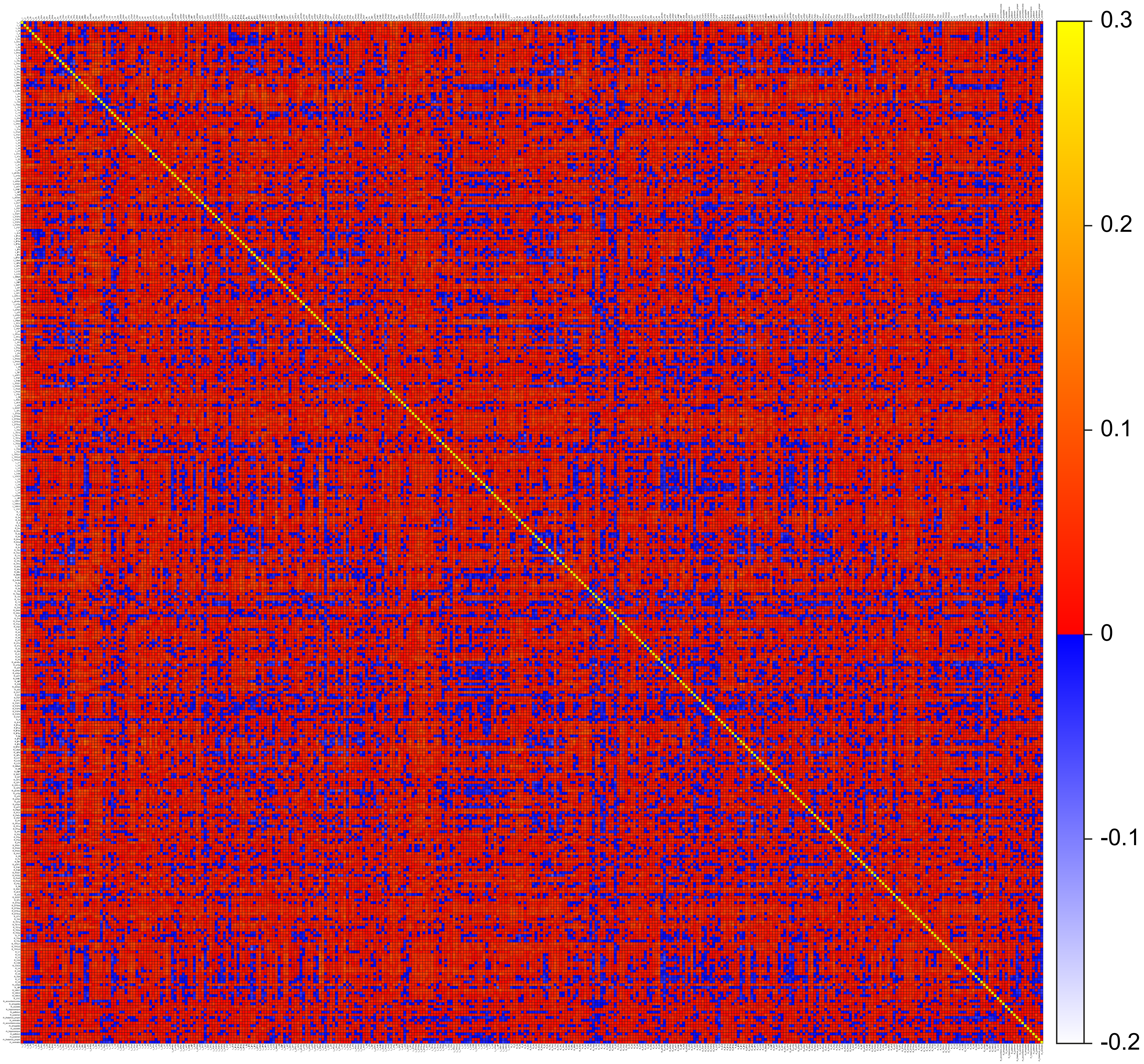

### Supplementary Figure 6

Changes in functional brain connectivity observed in the control group after fear acquisition training

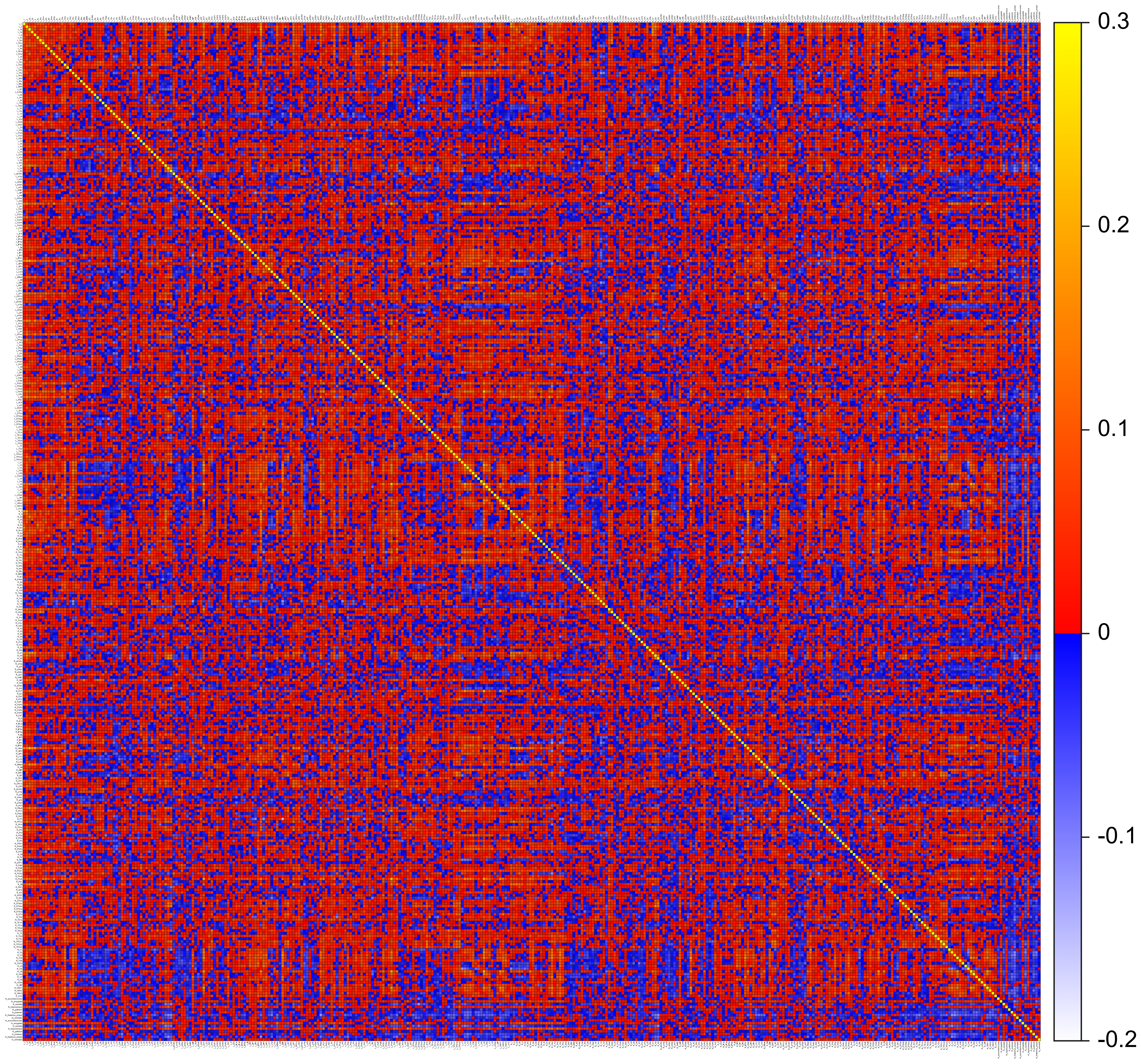

### Supplementary Figure 7

Significant brain activation during fear acquisition training (axial view)

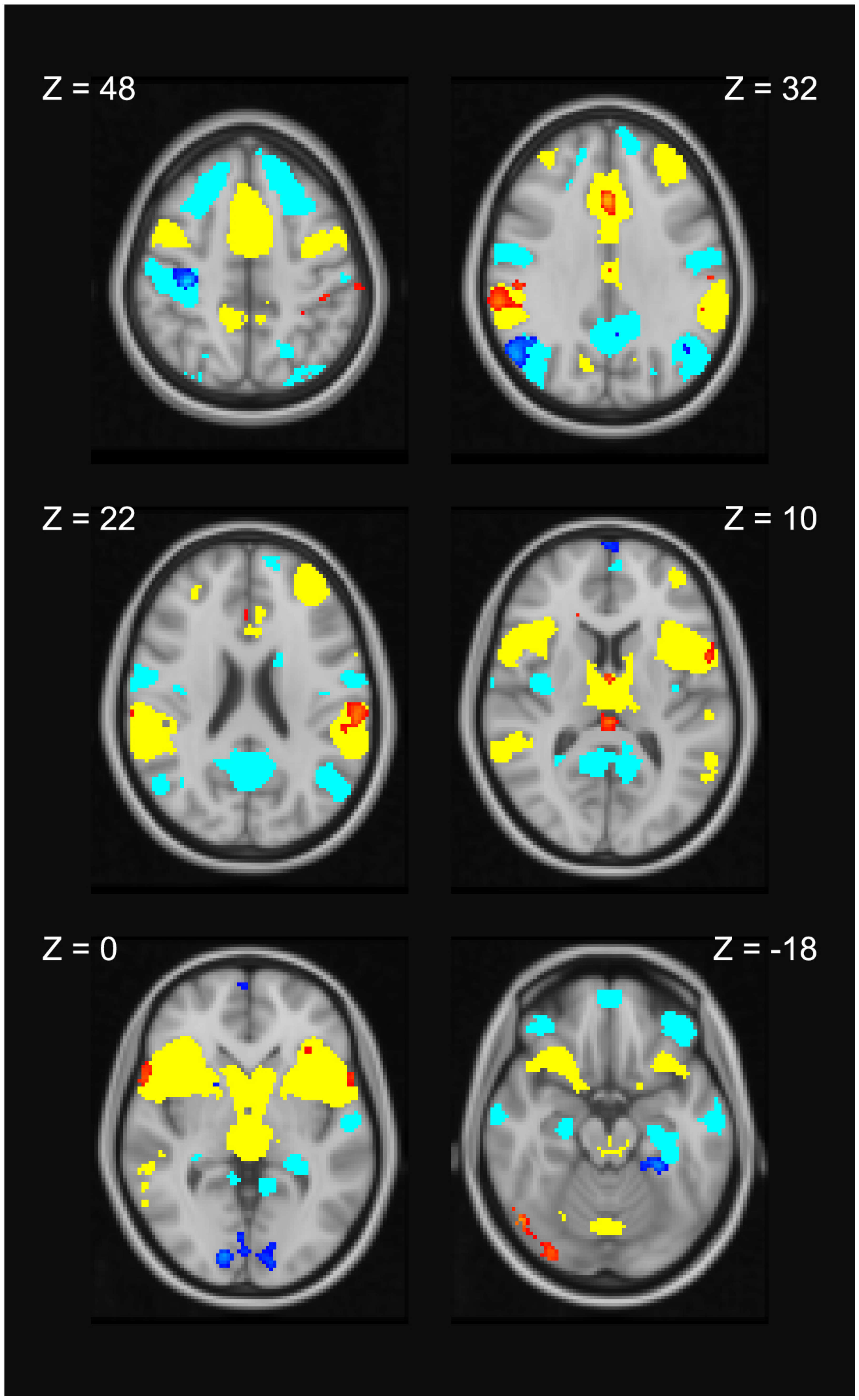

### Supplementary Figure 8

Significant brain activation during fear acquisition training (sagittal view)

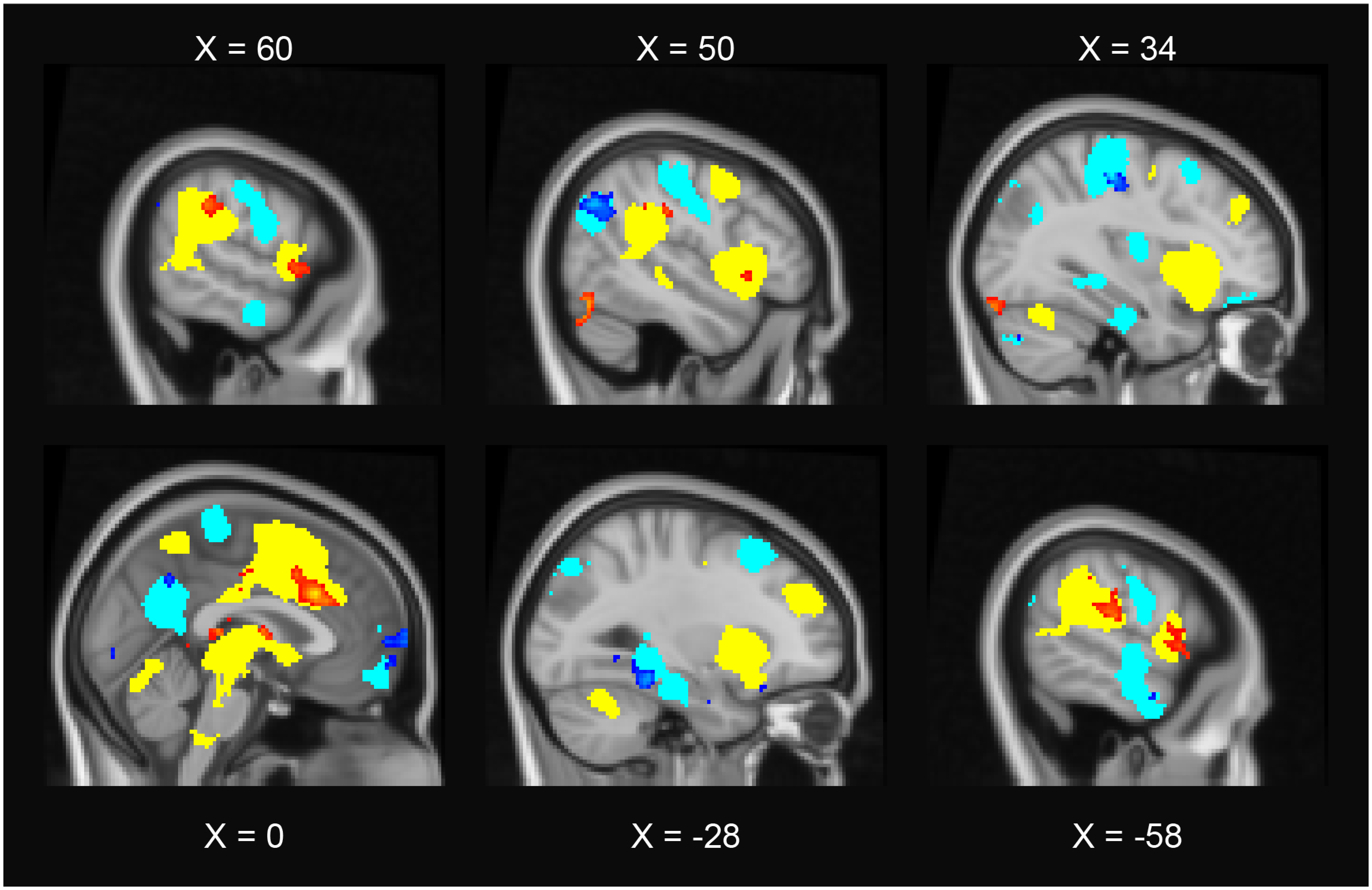

### Supplementary Figure 9

Partial regression diagrams showing the relationship between resting-state functional connectivity changes and mean conditioned fear responses

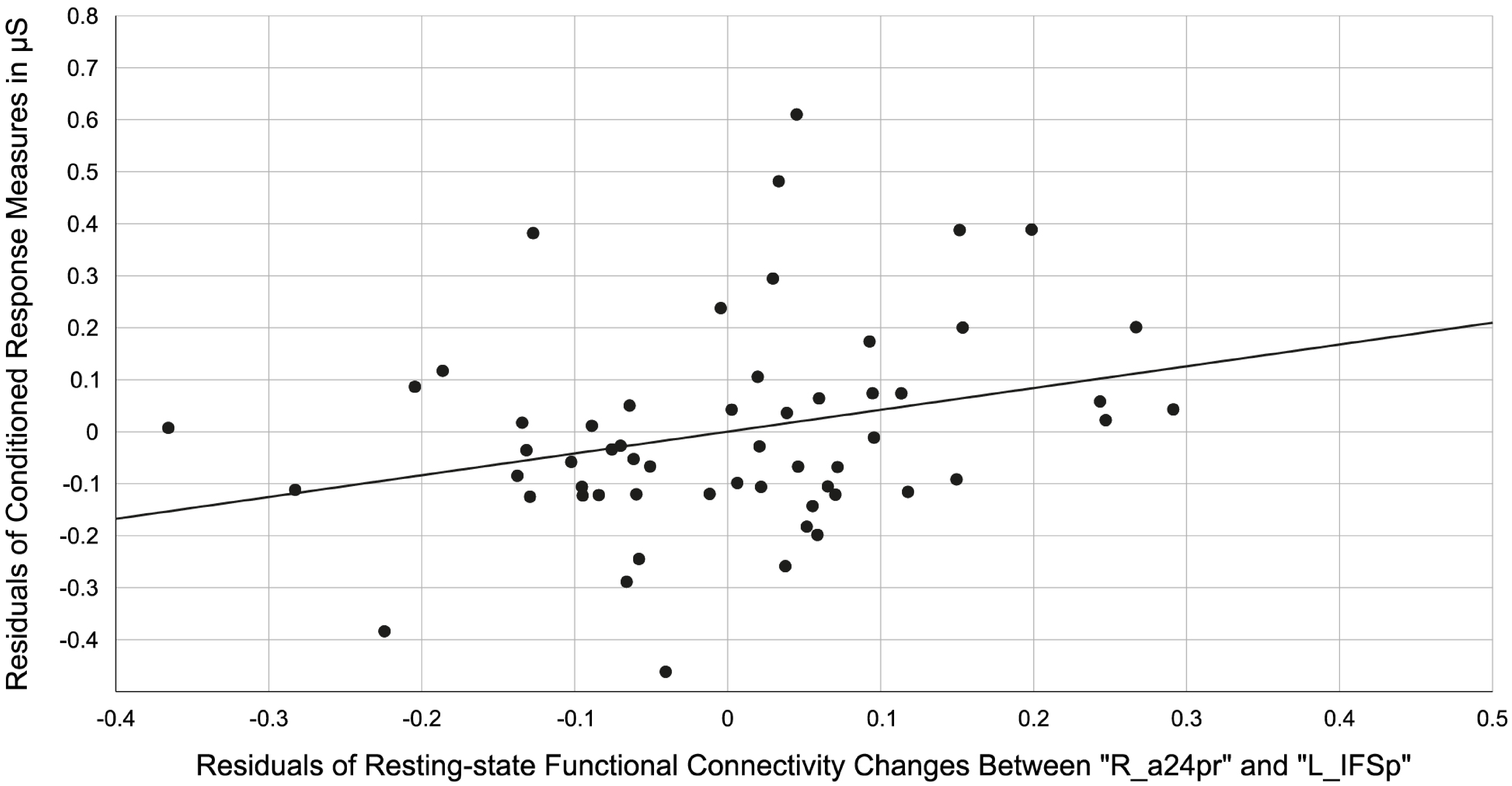


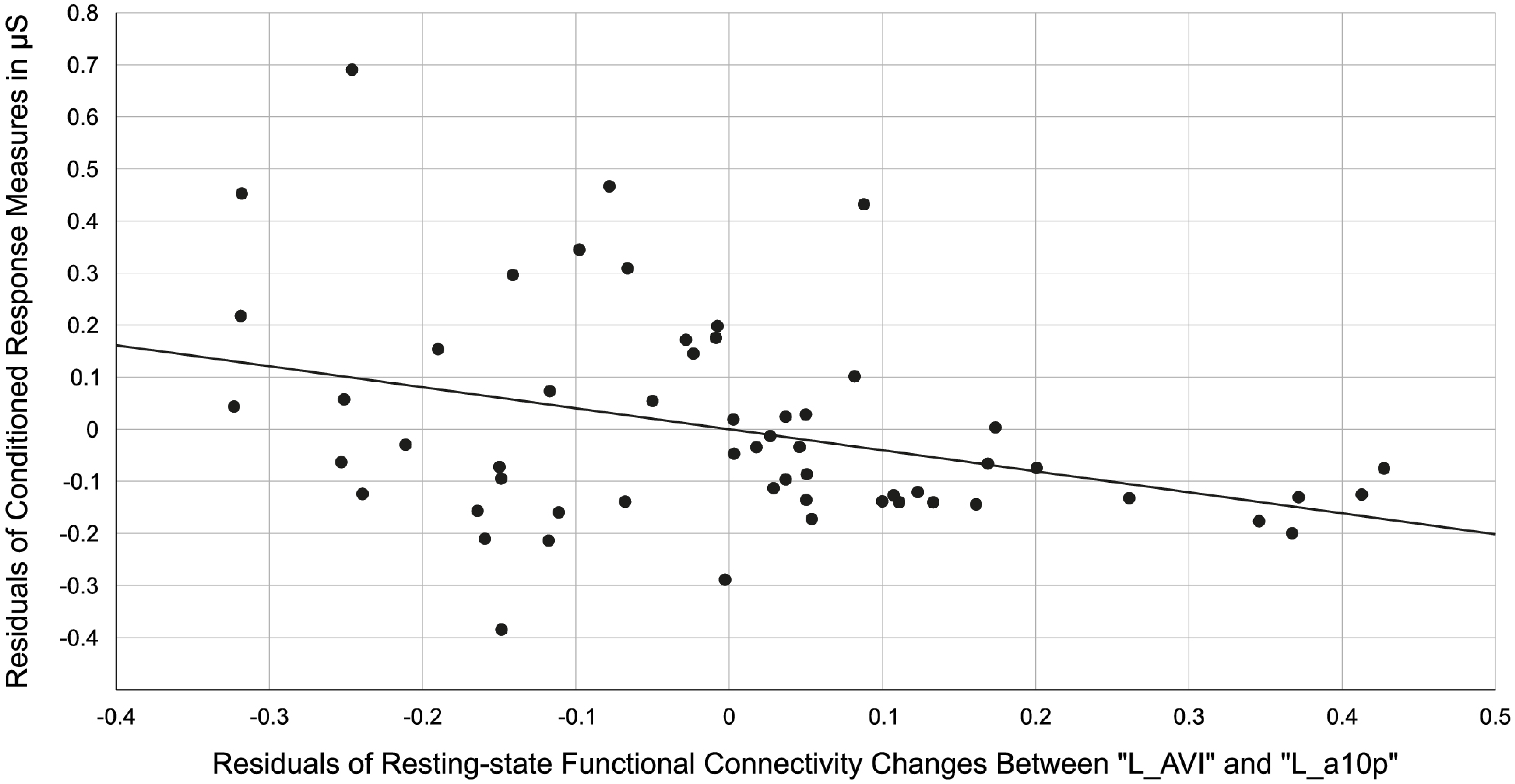

### Supplementary Figure 10

Connection-specific results from all three steps of the main analysis

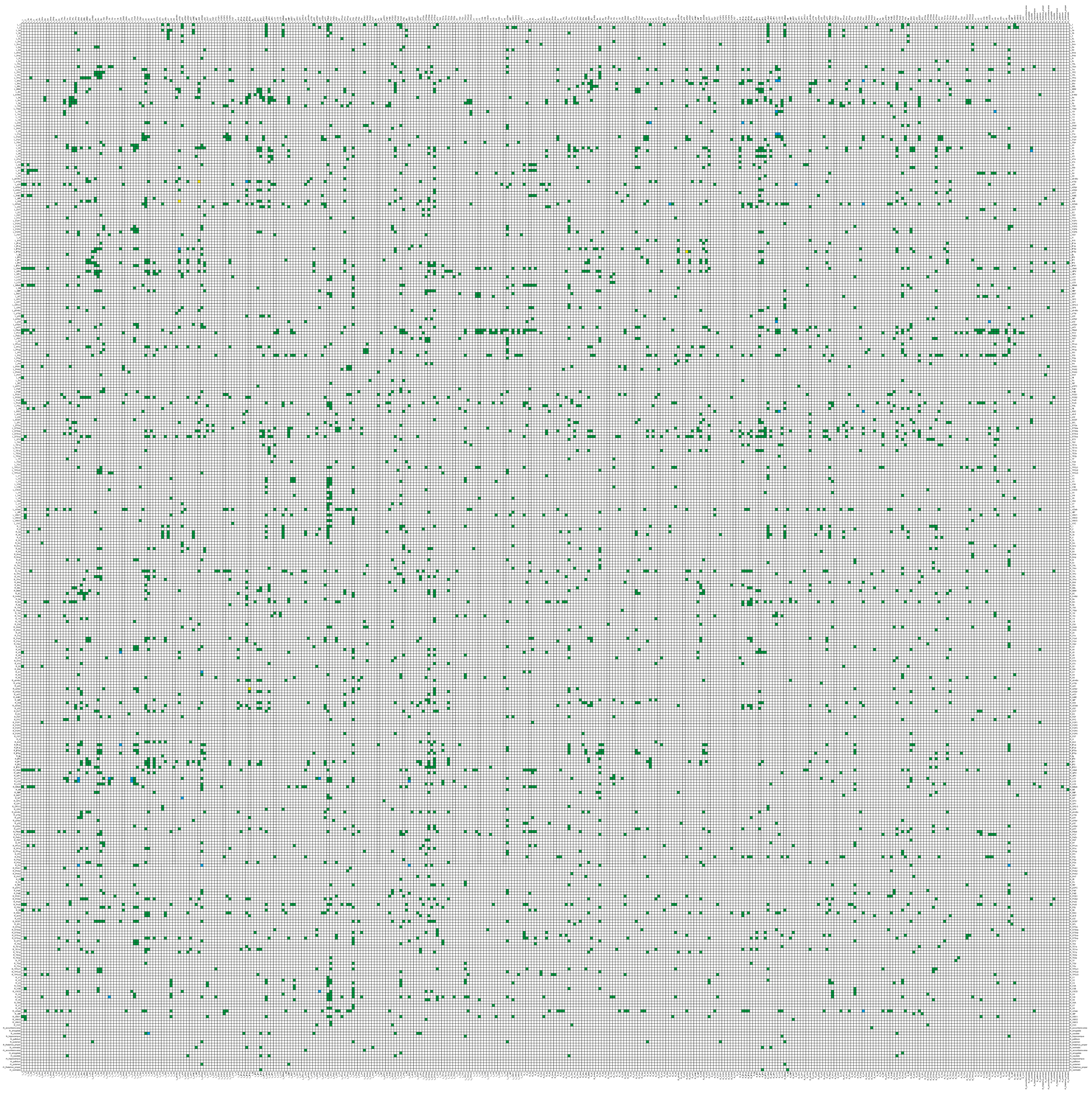
