## Supplementary Figure Captions for "Fear learning sculpts functional brain connectivity at rest beyond the traditional fear network in humans"

Supplementary Figure 1 - **Functional brain connectivity assessed in the experimental group prior to fear acquisition training.** Each cell in the symmetrical 374-by-374 matrix represents the functional connectivity between a pair of brain areas quantified as a Fisher *z*-transformed BOLD signal correlation. Data were assessed in the experimental group (*N* = 60) prior to fear acquisition training and controlled for head motion as well as signal from ventricles and white matter. Positive coefficients are displayed in red to yellow colors, while negative coefficients are displayed in blue to white colors. Auto-correlations on the diagonal are all displayed in yellow. Naming of brain areas is based on the Human Connectome Project's Multimodal Parcellation as well as FreeSurfer's Aseg Atlas.

Supplementary Figure 2 - **Functional brain connectivity assessed in the control group prior to fear acquisition training.** Each cell in the symmetrical 374-by-374 matrix represents the functional connectivity between a pair of brain areas quantified as a Fisher *z*-transformed BOLD signal correlation. Data were assessed in the control group (*N* = 24) prior to fear acquisition training and controlled for head motion as well as signal from ventricles and white matter. Positive coefficients are displayed in red to yellow colors, while negative coefficients are displayed in blue to white colors. Auto-correlations on the diagonal are all displayed in yellow. Naming of brain areas is based on the Human Connectome Project's Multimodal Parcellation as well as FreeSurfer's Aseg Atlas.

Supplementary Figure 3 - **Functional brain connectivity assessed in the experimental group after fear acquisition training.** Each cell in the symmetrical 374-by-374 matrix represents the functional connectivity between a pair of brain areas quantified as a Fisher *z*-transformed BOLD signal correlation. Data were assessed in the experimental group (*N* = 60) after fear acquisition training and controlled for head motion as well as signal from ventricles and white matter. Positive coefficients are displayed in red to yellow colors, while negative coefficients are displayed in blue to white colors. Auto-correlations on the diagonal are all displayed in yellow. Naming of brain areas is based on the Human Connectome Project's Multimodal Parcellation as well as FreeSurfer's Aseg Atlas.

Supplementary Figure 4 - **Functional brain connectivity assessed in the control group after to fear acquisition training.** Each cell in the symmetrical 374-by-374 matrix represents the functional connectivity between a pair of brain areas quantified as a Fisher *z*-transformed BOLD signal correlation. Data were assessed in the control group (*N* = 24) after fear acquisition training and controlled for head motion as well as signal from ventricles and white matter. Positive coefficients are displayed in red to yellow colors, while negative coefficients are displayed in blue to white colors. Auto-correlations on the diagonal are all displayed in yellow. Naming of brain areas is based on the Human Connectome Project's Multimodal Parcellation as well as FreeSurfer's Aseg Atlas.

Supplementary Figure 5 - **Changes in functional brain connectivity observed in the experimental group after fear acquisition training.** The value of each cell in the symmetrical 374-by-374 matrix was computed by subtracting the functional connectivity matrix assessed prior to fear acquisition training from the functional connectivity matrix assessed after fear acquisition training. Data were taken from the experimental group (*N* = 60) exclusively.

Positive values represent a shift towards more positive correlation coefficients, namely positive coefficients becoming larger, negative coefficients becoming smaller, or negative coefficients becoming positive. Negative values represent a shift towards more negative correlation coefficients, namely positive coefficients becoming smaller, negative coefficients becoming larger, or positive coefficients becoming negative. Positive values are displayed in red to yellow colors, while negative values are displayed in blue to white colors. Values on the diagonal are all displayed in yellow. Naming of brain areas is based on the Human Connectome Project's Multimodal Parcellation as well as FreeSurfer's Aseg Atlas.

Supplementary Figure 6 - **Changes in functional brain connectivity observed in the control group after fear acquisition training.** The value of each cell in the symmetrical 374-by-374 matrix was computed by subtracting the functional connectivity matrix assessed prior to fear acquisition training from the functional connectivity matrix assessed after fear acquisition training. Data were taken from the control group ($N$ = 24) exclusively. Positive values represent a shift towards more positive correlation coefficients, namely positive coefficients becoming larger, negative coefficients becoming smaller, or negative coefficients becoming positive. Negative values represent a shift towards more negative correlation coefficients, namely positive coefficients becoming smaller, negative coefficients becoming larger or positive coefficients becoming negative. Positive values are displayed in red to yellow colors, while negative values are displayed in blue to white colors. Values on the diagonal are all displayed in yellow. Naming of brain areas is based on the Human Connectome Project's Multimodal Parcellation as well as FreeSurfer's Aseg Atlas.

Supplementary Figure 7 - **Significant brain activation during fear acquisition training (axial view).** Voxels demonstrating significant BOLD signal contrasts between CS+ and CS- ($p < .05$, $Z > 3.1$, FWE-corrected) are displayed on a selection of axial slices from the Montreal Neurological Institute 152 T1 2-mm template. Brain activation in response to the CS+ versus the CS- is depicted in red to yellow colors, while brain activation in response to the CS- versus the CS+ is depicted in blue to light blue colors. Uniformly colored voxel clusters represent CS+ > CS- activation (yellow) and CS- > CS+ activation (light blue) as reported in a meta-analysis of fMRI studies on fear conditioning (Fullana MA et al. 2016).

Supplementary Figure 8 - **Significant brain activation during fear acquisition training (sagittal view).** Voxels demonstrating significant BOLD signal contrasts between CS+ and CS- ($p < .05$, $Z > 3.1$, FWE-corrected) are displayed on a selection of sagittal slices from the Montreal Neurological Institute 152 T1 2-mm template. Brain activation in response to the CS+ versus the CS- is depicted in red to yellow colors, while brain activation in response to the CS- versus the CS+ is depicted in blue to light blue colors. Uniformly colored voxel clusters represent CS+ > CS- activation (yellow) and CS- > CS+ activation (light blue) as reported in a meta-analysis of fMRI studies on fear conditioning (Fullana MA et al. 2016).

Supplementary Figure 9 - **Partial regression diagrams showing the relationship between resting-state functional connectivity changes and mean conditioned fear responses.** Scatter plots illustrate the results yielded by multiple regression analysis with backward elimination. For each participant, changes in resting-state functional connectivity were computed by subtracting the functional connectivity matrix assessed prior to fear acquisition training from the functional connectivity matrix assessed after fear acquisition training. Increases in resting-state functional connectivity are indicated by positive values and decreases by negative values. Mean conditioned fear responses were computed by subtracting the average skin conductance responses across all CS- trials from the average skin conductance responses across all CS+ trials.

Supplementary Figure 10 - **Connection-specific results from all three steps of the main analysis.** Each cell in the symmetrical 374-by-374 matrix represents a functional connection between a pair of brain areas and is either depicted in green, light blue, yellow, or white. Green color indicates that the respective connection showed a statistically significant increase or decrease due to fear acquisition training, i.e. passing the first step of the main analysis. Light blue color indicates that the respective connection also exhibited a statistically significant result in the comparison of functional connectivity changes between the experimental and control group, i.e. passing the first two steps of the main analysis. Yellow color indicates that the respective connection's change in functional connectivity was also associated with conditioned fear response measures quantified via skin conductance responses, i.e. passing all three steps of the main analysis. White color indicates that the respective connection did not show any significant changes due to fear acquisition training and was not subjected to any other steps of the main analysis. Cells on the diagonal represent auto-connections and are all displayed in white. The first two steps of the analysis included a correction for multiple comparisons using the Benjamini-Hochberg method. Naming of brain areas is based on the Human Connectome Project's Multimodal Parcellation as well as FreeSurfer's Aseg Atlas.
