## Supplementary Table 1 for "Fear learning sculpts functional brain connectivity at rest beyond the traditional fear network in humans"

Supplementary Table 1. Significant r*esults yielded by the comparison of difference matrices from the experimental group and the control group*

| **Area A** | **Area B** | ***t*-value** | ***p*-value** |
|---|---|---|---|
| R_PH (49/-60/-11) | L_8Ad (-23/28/43) | 4.754 | .000008 |
| R_LO2 (45/-80/-5) | L_31a (-5/-35/43) | 4.421 | .000030 |
| R_LO1 (40/-81/3) | L_p32 (-11/48/0) | 4.374 | .000036 |
| R_LO1 (40/-81/3) | L_10r (-7/52/-7) | 4.315 | .000044 |
| R_IFJa (41/20/23) | L_23d (-2/-18/38) | 4.289 | .000049 |
| R_LO1 (40/-81/3) | L_8Ad (-23/28/43) | 4.163 | .000077 |
| R_a24pr (5/20/33)* | L_IFSp (-47/23/21)* | 4.091 | .000100 |
| L_IFSa (-45/33/12) | L_a10p (-25/58/-5) | 4.083 | .000103 |
| lh_caudate (-13/7/11) | L_45 (-48/28/5) | 4.016 | .000130 |
| R_V4t (46/-76/-2) | L_10r (-7/52/-7) | 3.949 | .000165 |
| R_PH (49/-60/-11) | L_RSC (-5/-38/18) | 3.863 | .000223 |
| R_A1 (44/-21/10) | L_d23ab (-2/-38/31) | 3.858 | .000226 |
| R_PH (49/-60/-11) | L_d23ab (-2/-38/31) | 3.833 | .000247 |
| R_LO2 (45/-80/-5) | L_8Ad (-23/28/43) | 3.792 | .000285 |
| R_V3CD (35/-81/14) | L_p32 (-11/48/0) | 3.747 | .000332 |
| L_AVI (-31/25/-2)* | L_a10p (-25/58/-5)* | 3.715 | .000370 |
| R_v23ab (6/-51/19) | R_PH (49/-60/-11) | 3.713 | .000372 |
| R_45 (50/29/4) | L_23d (-2/-18/38) | 3.680 | .000416 |
| R_LO2 (45/-80/-5) | L_RSC (-5/-38/18) | 3.671 | .000429 |
| R_LO1 (40/-81/3) | L_31a (-5/-35/43) | 3.650 | .000460 |
| R_MT (48/-70/6) | L_a24 (-5/41/0) | 3.628 | .000496 |

Twenty-one functional connections which exhibited significant differences in their resting-state functional connectivity changes between the experimental and control groups according to two-sample t-tests ($p < .05$, Benjamini-Hochberg corrected). Functional connections are constituted by areas A and B. Naming of brain areas is based on the Human Connectome Project's Multimodal Parcellation as well as FreeSurfer's Aseg Atlas. MNI coordinates of an area's center of gravity are given in brackets. Asterisks highlight functional connections in which resting-state functional connectivity changes were significantly associated with the conditioned reaction of participants as quantified via skin conductance responses.
